# Integrative Multiomic Analysis Reveals How Non-Viral Delivery System Selection Shapes CRISPR Gene Editing Outcomes in Stem Cells

**DOI:** 10.64898/2026.05.29.728905

**Authors:** Joshua P. Graham, Alexander V. Arteaga, Abolfazl S. Moghaddam, Kara L. Spiller, Daniel J. Laverty, Tomas Gonzalez-Fernandez

## Abstract

The clinical translation of CRISPR gene editing is challenged by the lack of delivery systems that are both safe and efficient in therapeutically relevant cell types such as mesenchymal stem cells (MSCs). Non-viral delivery avoids the immunogenicity and genomic integration risks of viral vectors but faces fundamental trade-offs between editing efficiency and cytotoxicity. Here, we present a comprehensive multiomic analysis of four non-viral CRISPR delivery modalities including cell-penetrating peptide- (CPP), lipid-, and polymer-based nanoparticles and electroporation; across mRNA and ribonucleoprotein (RNP) molecular formats. We systematically evaluate each modality, demonstrating that lipid-based delivery achieved the highest editing rates at the cost of genomic instability risks, interferon pathway activation, and a pro-inflammatory shift in MSC paracrine activity. Alternatively, CPPs yield moderate editing rates while reducing these unintended side-effects, whereas polymers and electroporation consistently yielded the lowest efficiencies. CRISPR molecular format and delivery method interacted in a stress-dependent manner, with RNP delivery reducing editing rates under high-stress systems while improving them in lower-stress modalities such as CPP and electroporation. These findings establish that editing efficiency alone is an insufficient metric for delivery system selection, and that genomic stability, transcriptomic dysregulation, and inflammatory response must be treated as primary design criteria for CRISPR therapies.

## 1. Introduction

Clustered regularly interspaced short palindromic repeats (CRISPR) gene editing has revolutionized biomedicine by adding unprecedented genomic and transcriptomic control at specific loci.^1^ This highly precise and tunable platform has been employed for knock-out screens,^2–4^ transgene insertion,^5–7^ epigenetic regulation,^8,9^ and prime/base editing^10–12^ to better understand biological pathways, correct genetic disorders, and alter cell phenotypes. Yet the clinical application of these strategies remains limited by the difficulty of efficiently delivering all the necessary components to therapeutically relevant cell types.

The most common CRISPR delivery strategy involves packaging CRISPR associated (Cas) nucleases and guide RNAs (gRNAs) into viral vectors for reliable, long-term expression after transduction. But the inherent immunogenicity of viral vectors continues to cause adverse effects that negatively impact therapeutic outcomes.^13–15^ Furthermore, while Cas9 itself fits adeno-associated virus (AAV) packaging limits, Cas9 fusion proteins such as those used for CRISPR activation and inhibition require the transduction and splicing of multiple vectors which reduces efficiencies and burdens biomanufacturing pipelines.^16^ This complexity also increases risks of AAV concatemerization and host genome integration which have been implicated in liver fibrosis and necrosis in humans treated with AAV gene therapies.^17^ These risks were recently demonstrated clinically in a patient that suffered a neuroepithelial tumor which was attributed to AAV integration following an AAV serotype 9 gene therapy.^18^ Alternatively, lentiviral vectors offer higher packing limits, but the risk of insertional mutagenesis greatly increases off-target editing and genomic instability risks.^19^ These challenges highlight the need for non-viral gene editing strategies for effective *in vivo* and *ex vivo* gene editing.

Non-viral CRISPR strategies employ physical methods such as electroporation or nanoparticle-based systems to deliver CRISPR machinery in plasmid DNA (pDNA), mRNA, or ribonucleoprotein (RNP) molecular formats.^20^ Interest in non-viral delivery has grown considerably following the FDA approval of mRNA-based COVID-19 vaccines, which demonstrated that lipid nanoparticles (LNPs) could deliver nucleic acid cargo with high efficiency at a clinical scale.^21,22^ Yet, LNPs are inherently immunogenic, helping to mount a response to the delivered spike protein and initiate antibody production. Pongma *et al.* demonstrated that LNPs delivered via intramuscular injection activated inflammatory IL-6/JAK/STAT3 and TNFα/NF-κB pathways in resident liver macrophages and polarized them towards a pro-inflammatory phenotype.^23^ This immune activation limits the clinical use of LNPs for CRISPR delivery, as both Cas proteins and gRNAs individually trigger inflammatory pathways.^24^ In fact, Li *et al*. demonstrate that pre-existing Cas9 immunity in mice causes a cytotoxic t-cell response that eliminates gene editing outcomes.^25^ This is especially concerning since antibodies against *Staphylococcus aureus* and *Streptococcus pyogenes* Cas9 have been detected in 78% and 58% of human donors respectively.^26^ Furthermore, 5’ triphosphate groups generated during *in vitro* transcription of gRNAs activate the retinoic acid-inducible gene I (*RIGI*) pathway and downstream cell stress and apoptosis pathways.^24^ Therefore, there is a strong need to minimizing immune activation during CRISPR delivery.

Beyond immune activation, the unintended transcriptomic dysregulation resulting from transfection can also confound the interpretation of gene editing outcomes. CRISPR screens are engineered to characterize genes through highly specific editing but disrupting the expression of hundreds to thousands of genes during CRISPR delivery could conceal the functional impacts of specific gene edits and increase false positive rates. For example, LNP delivery to T cells induces thousands of differentially expressed genes (DEGs).^27^ Similarly, other lipid-based systems have demonstrated widespread transcriptomic dysregulation and inflammatory signaling.^28,29^ Efforts must be taken to minimize these unintended transcriptomic effects for the accurate interpretation of gene editing outcomes and designing reproduceable strategies.

Transcriptomic dysregulation and the inflammatory effects of transfection are especially concerning in stem cell therapies. Weinert *et al.* reported that stem cells were significantly more sensitive to the inflammatory effects of *in vitro* transcribed gRNAs compared to cell lines.^24^ Human mesenchymal stem cells (MSCs) are among the most widely used cell types in clinical trials, where their efficacy in regenerative medicine and immunotherapy relies on multipotency, proliferation, self-renewal, and paracrine secretory properties. Therefore, preserving these essential stem cell characteristics is critical when designing CRISPR strategies for MSC engineering.^30^ In fact, the choice of transfection reagent has been shown to significantly alter the viability and differentiation of primary MSCs.^30^ Careful design of CRISPR delivery strategies is essential for maximizing editing rates while maintaining cell health and proliferation to achieve the highest therapeutic efficacy.

The objective of this study is to systematically compare non-viral systems for the delivery of CRISPR cargos, characterizing the functional and phenotypic trade-offs of each strategy. To accomplish this, we first establish cell penetrating peptide (CPP), lipid, and polymeric nanoparticle or electroporation systems by analyzing the fluorescent profiles following eGFP mRNA delivery then compare these results to the editing efficiencies from each method. Next, we performed bulk RNA sequencing and data-dependent acquisition (DDA) proteomics to characterize the transcriptomic dysregulation induced by each delivery approach. Then, we assessed the downstream impact on MSC immunomodulatory function through co-culture with primary human monocyte-derived macrophages (MDMs). Finally, we compare the transcriptomic signatures of CRISPR delivery in mRNA and RNP molecular formats. Our results demonstrate that non-viral CRISPR delivery in MSCs features many trade-offs between editing efficiency, genomic instability, cytotoxicity, and immunomodulation which must be considered when designing CRISPR and other gene therapy-based stem cell therapies.

## 2. Materials and Methods

### 2.1. Materials

pX458-Ef1a-Cas9-H2B-mCherry was a gift from Joseph Brzezinski (Addgene plasmid # 159655; http://n2t.net/addgene:159655; RRID:Addgene_159655).^31^ EZ Cap™ Cas9 mRNA (m1Ψ) was purchased from APExBIO. TrueCut™ HiFi Cas9 Protein was purchased from Thermo Fisher. The gRNA sequence targeting the human *AASV1* locus (GGGGCCACTAGGGACAGGAT) was identified previously.^32^ Oligonucleotides for the production of gRNAs were produced by Thermo Fisher. gRNAs were cloned into Cas9-mCherry pDNA by phosphorylating and annealing the primers using the T4 polynucleotide kinase kit (NEB), digesting addgene 159655 with *BbsI* (NEB), and annealing these products using the Quick Ligase (NEB). gRNAs for co-delivery with mRNA or RNP complexing were produced using the precision gRNA synthesis kit (Thermo Fisher).

The RALA peptide (WEARLARALARALARHLARALARALRACEA) was produced by chemical peptide synthesis at Genscript, USA and provided as a lyophilized powder at 96.6% purity via high-performance liquid chromatography (HPLC). Lipofectamine MessengerMAX and Lipofectamine CRISPRMAX were purchased from Thermo Fisher. Branched 25 kD polyethyleneimine (PEI) was purchased from Millipore Sigma.

### 2.2 Cell culture and transfections

#### 2.2.1 Cell culture

Human bone marrow derived mesenchymal stem cells (hMSCs) were obtained from RoosterBio (donor 310272). Cells were grown in expansion media (DMEM supplemented with 10% fetal bovine serum (FBS) (Gemini Bio) 1% penicillin/streptomycin (Thermo Fisher)) with media changes every 2-3 days until confluency. To passage cells, they were first lifted with trypsin (0.25% trypsin, 0.1% disodium ethylene diaminetetraacetic acid (EDTA) in Hanks’ balanced salt solution (HBSS) (Corning) for 5 minutes. Trypsin was then inactivated with expansion media. Cells were then pelleted by centrifugation for 5 minutes at 300 xg, resuspended in fresh expansion media and reseeded at 5.7×10^3^ cells/cm^2^ on tissue culture plastic treated for cell adhesion.

Human buffy coats were obtained from the American National Red Cross within one day of blood donation and monocytes were isolated as previously reported.^33^ Buffy coats were diluted 2x in 1 mM EDTA in PBS then 20 mL was slowly laid over 15 mL of Ficoll-Paque Plus reagent (Cytiva) and centrifuged at 400 xg for 20 minutes at room temperature. The white band of peripheral blood mononuclear cells (PBMCs) was removed using a sterile transfer pipette. PBMCs were washed twice by diluting or resuspending in 50 mL in 1 mM EDTA in PBS then pelleting at 150 xg for 10 minutes. PBMCs were resuspended in in RPMI 1640 media (Gibco) supplemented with 10% FBS, 1% pen/strep, counted using the Countess III automated cell counter (Invitrogen), and diluted to 2 × 10^6^ cells/mL. 25 mL of the diluted PBMCs were gently overlaid on 25 mL of Percoll solution (48.6 % RPMI, 5.4% FBS, 0.345 x PBS, and 42.55% Percoll Plus reagent (Cytiva)) and centrifuged at 550 xg for 30 minutes.

The white band of monocytes at the interface of the upper and lower solutions was collected and washed in 1 mM EDTA in PBS by centrifuging at 400 xg for 10 minutes. Finally, the cells were counted in PBS and frozen in RPMI supplemented with 10% FBS, 1% pen/strep, and 5% dimethyl sulfoxide (DMSO).

#### 2.2.2 Transfections

RALA transfections were performed as previously reported.^34^ Confluent MSCs were lifted with trypsin for 3 minutes followed by inactivation with expansion media. Cells were pelleted by centrifugation at 300 xg for 5 minutes, washed with optiMEM (Gibco), and finally resuspended in OptiMEM at 1 × 10^6^ cells/mL. 1 × 10^4^ cells per cm^2^ of culture area were mixed with complexed RALA-CRISPR nanoparticles. For RNP delivery, 4 × 10^2^ ng/cm^2^ of Cas9 was mixed with a 1:1 molar ratio of *AAVS1* guide RNA (gRNA) (GGGGCCACTAGGGACAGGAT) and incubated for 15 minutes before complexing with RALA at 150x RALA:RNP molar ratio. Finally, for RNA delivery, 1.3 × 10^2^ ng/cm^2^ of mRNA at a 1:10 molar ratio of mRNA:gRNA was complexed with RALA at an NP ratio of 7. Nanoparticles were incubated with the cells for 10 minutes at 37 °C before cells were moved to 6-well plates and allowed to adhere. After 6 hours, OptiMEM was aspirated, and cells were washed with PBS then cultured in 2 mL of expansion media.

Lipofectamine was used according to manufacturer protocols. Briefly, hMSCs were seeded at 2.6 × 10^3^ cells/cm^2^ in 6 well plates 24 hours before transfection. 1.3 × 10^2^ ng/cm^2^ of mRNA/gRNA was directly mixed with the 3 μL Lipofectamine MessengerMAX reagent (Invitrogen) per μg of mRNA, incubated for 5 minutes, and added to cells. 2 × 10^2^ ng/cm^2^ of Cas9 was mixed with a 1:1 molar ratio of gRNA and incubated for 15 minutes. 4 × 10^−3^ μL/cm^2^ of Cas9^+^ reagent in 13 μL/cm^2^ of OptiMEM. This RNP solution was then combined 1:1 with 2.4 ×10^−3^ μL/cm^2^ of Lipofectamine CRISPRmax in 13 μL/cm^2^ of OptiMEM. 13 μL/cm^2^ was then added directly to the adhered cells.

PEI transfections were performed as previously reported.^30,34^ Briefly, cells were seeded at 2.6 ×10^3^ cells/cm^2^ in 6-well plates 24 hours before transfection then washed with PBS and incubated in OptiMEM 2 hours before transfection. mRNA nanoparticles were prepared by mixing 1.3 × 10^2^ ng/cm^2^ of total RNA with PEI at NP 7. RNP nanoparticles were made by mixing 2 × 10^2^ ng/cm^2^ of Cas9 with gRNA at a 1:1 molar ratio and incubating for 15 minutes before adding 5 μL/cm^2^ of water and 1.39 × 10^−1^ μg/cm^2^ of PEI. These particles were diluted 10:1 vol/vol in OptiMEM and 500 μL of this mixture was added to each well then diluted with 1 mL of OptiMEM after 15 minutes. Finally, OptiMEM was removed, and cells were washed with PBS and 2 mL of expansion media was added to each well 4 hours after transfection.

#### 2.2.3. Electroporation

Cells were lifted using trypsin, diluted in XPAN, washed in optiMEM, then resuspended in Gene Pulser electroporation buffer (Bio-Rad) at 1 × 10^6^ cells/mL. 400 μL of cells were then mixed with 10μg of pDNA, 10 μg of RNA (10:1 moles gRNA:mRNA), or 10 μg of complexed RNP, incubated on ice for 15 minutes, then electroporated at 150 V, 1000 μF and 200 Ω as optimized (Fig. S1).^35^ The were immediately added to warm expansion media to recover.

### 2.3 Sanger sequencing to analyze Cas9 editing efficiencies

Total genomic DNA was extracted from transfected-or non-transfected cells using the Monarch Spin gDNA Extraction Kit (NEB). Primers were designed using NCBI primer-blast (https://www.ncbi.nlm.nih.gov/tools/primer-blast/). A 1000 bp region centered at the *AAVS1* cut site targeted by our gRNA was analyzed against the Refseq representative genomes database yielding the forward primer TCCTCTCTGGCTCCATCGTA and reverse primer CCCGTTCTCCTGTGGATTCG which had minimal off-target binding sites compared to the remaining genome. Primers were produced by Thermo Fisher with purification by desalting. PCR amplification was performed using the NEBNext® Q5® Hot Start HiFi PCR Master Mix (NEB) with 0.3 mM primers and 10 ng of genomic DNA (Fig. S2). The thermocycler conditions are described in table S1. Sanger sequencing of the PCR products was performed by Genewiz and the resulting .ab1 files were input into Deconvolution of Complex DNA Repair (DECODR, https://decodr.org/) to determine the size and abundance of indels following gene editing.

### 2.4 Profiling the transfection of MSCs

Flow cytometric analysis of MSCs transfected with GFP mRNA was performed as previously reported.^34^ Briefly, MSCs were lifted using trypsin, diluted in expansion media, centrifuged for 5 minutes at 500 xg, and resuspended in PBS. Samples were analyzed using a cytoflex flow cytometer (Beckman Coulter, USA). The forward scatter (FSC), side scatter (SSC), and FITC channel gains were set to 20, 20, and 1 respectively.

Cell viability following transfection was determined using the alamarBlue reagent (Invitrogen) according to the manufacturer. Media was removed completely from the cells and replaced with alamarBlue reagent diluted 1:10 in expansion media. The cells were incubated at 37°C and 5% CO_2_ for 45 minutes, then the alamarBlue solution was collected and 3 technical replicates were analyzed at an excitation of 550 nm and emission at 590 nm on a SpectraMax ID3 plate reader (Molecular Devices).

### 2.5 mRNA sequencing

Total RNA was extracted on 2 days after transfection using the Trizol reagent (Invitrogen) then purified using PureLink™ RNA Mini Kit (Invitrogen) according to the manufacturers protocol. Total RNA was sequenced by Novogene using the NovaSeq X Plus Series platform (PE150) with 6 G of raw data per sample. The resulting .fastq files were processed using kallisto with mapping to the Ensembl human reference transcriptome (GRCh38, release 111, cDNA, all transcripts) yielding estimated counts and transcripts per million (tpm)( https://useast.ensembl.org/Homo_sapiens/Info/Index).^36^ All RNA-seq data from this experiment is available on the Gene Expression Omnibus (GEO) at accession GSE333489. This output was then normalized using sleuth, kallisto’s complement for normalizing and analyzing data, aggregating by gene name and using the log2(x + 0.5) normalization function (https://hbctraining.github.io/main). Sleuth was also used for differential expression analysis with reference to non-transfected control samples.

Normalized RNAseq counts and differential expression data were then processed using python 3.12.2. The global transcriptomic profile of the RNAseq counts was analyzed using uniform manifold approximation and projection (UMAP, v0.5.7) (n_components = 2, n_neighbors = 5) and t-SNE (perplexity = 6, 1000 iterations). UMAP results were also assessed after dimensionality reduction via principal component analysis (PCA)(Fig. S3). K-means clustering was performed using the scikit-learn implementation (sklearn.cluster.KMeans, v1.5.2) with the optimal number of clusters (k = 5) determined empirically by the elbow method, minimizing within-cluster inertia (Fig. S4). A fixed random seed (random_state = 23) was used to ensure reproducibility. K-means clustering results were visualized by projecting samples onto the first two principal components via PCA (sklearn.decomposition.PCA, n_components = 2), with samples colored by cluster assignment. Variance explained by PC1 and PC2 is reported on the respective axes.

To visualize group separation in UMAP, t-SNE, and PCA embedding spaces, confidence ellipses were computed for each treatment group and overlaid on each two-dimensional projection. For each group, the covariance matrix of the embedded coordinates was decomposed via eigenvalue analysis to determine the orientation and magnitude of variance. Ellipse geometry was defined by the principal eigenvectors and scaled to encompass 2 standard deviations along each axis, such that the ellipse width and height correspond to 2 × 2√λ where λ represents the eigenvalues of the covariance matrix. Ellipses were rendered using matplotlib’s Ellipse patch and are colored by treatment group. To assess global similarity, the Jensen–Shannon distance was calculated across all genes and the resulting distance matrix was used to generate a clustermap plot.

Volcano plots were produced using plotly.express by filtering differentially expressed genes (DEGs) for an adjusted p value (q) less than 0.05 and a log_2_ Fold Change (log_2_FC) greater than 1 or less than −1. Gene set enrichment analysis (GSEA) was performed using the gseapy package prerank function, with DEGs ranked according to **equation 1**.

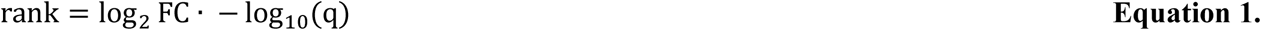

Ranked genes were analyzed with reference to the “MSigDB_Hallmark_2020” gene set (https://www.gsea-msigdb.org/gsea/msigdb/human/genesets.jsp?collection=H) with the parameters: min_size = 5, max_size = 1000, permutation_num = 1000. The resulting gene ontology terms were filtered for those with a false discovery rate (FDR) less than 0.25 and the normalized enrichment score (NES) of each term was plotted as a heat map using the seaborn package.^37^ Gene sets and their defined abbreviations are provided in table S2.

### 2.6 Bottom-Up Proteomics

#### 2.6.1 Proteomics Sample Preparation

MSCs (750,000 cells) were transected as described. After two days, the cells were lysed 50 mM Tris-HCl (pH 7.5), 150 mM NaCl, 1% (w/v) sodium dodecyl sulfate (SDS), and 0.1% (v/v) Triton X-100. Cell debris was removed by centrifugation at 12,000 × *g* for 12 minutes at 4°C and the protein concentration in the clarified supernatant was determined by the bicinchoninic acid (BCA) assay (Thermo Fisher). 100 μg of protein from each sample was diluted to attain a final concentration of 50 mM ammonium bicarbonate (ABC, pH 8) and 5% (w/v) SDS. Samples were reduced by adding tris(2-carboxyethyl)phosphine (TCEP) to 10 mM final concentration and incubating at 95°C for 3 minutes before cooling to room temperature. Cysteine residues were subsequently alkylated by addition of iodoacetamide (IAA) to a final concentration of 20 mM which was incubated for 30 minutes at room temperature in the dark.

Proteins were then captured on EZ-10 spin column silica membranes according to Miniprep-Assisted Proteomics (MAP) protocols.^38^ Ammonium acetate (5 M, pH 5.0) was added at a 1:10 (v/v) ratio, followed by seven volumes of 95:5 (v/v) methanol: 500 mM ABC (500 mM final concentration) mixed gently by pipetting. The suspension was loaded onto an EZ-10 spin column in a 2 mL collection tube and centrifuged at 1,200 × *g* for 30 sec; the flow-through was discarded. Bound proteins were washed twice with 500 µL of 95:5 (v/v) methanol:ABC (500 mM final concentration) at 1,200 × *g* for 30 seconds per wash to remove residual SDS, TCEP, IAA, and salts.

On-column trypsin digestion was performed at a 1:50 (w/w) enzyme:protein ratio in 50 μL volume. Columns were sealed in a humidified container to prevent evaporation and incubated at 37°C overnight. Peptides were eluted by sequential centrifugation at 1,200 × *g* for 60 seconds in 0.1% (v/v) formic acid in water and 0.2% (v/v) formic acid in 1:1 acetonitrile:water. Eluates were pooled and concentrated to approximately 50–100 µL by nitrogen gas purging. Samples were then desalted using pre-conditioned C18 solid-phase extraction (SPE) cartridges. Peptides were added to the column and washed three times with 0.1% formic acid in water before elution. Peptides were eluted stepwise with 2 mL each of 30%, 50%, 70%, 80%, and 20% acetonitrile in 0.1% formic acid. Eluates were dried to approximately 30 µL by nitrogen gas purging or vacuum centrifugation. Peptide concentration was assessed by UV absorbance (NanoDrop), and complete digestion was confirmed by SDS-polyacrylamide gel electrophoresis (SDS-PAGE). Data was collected immediately after sample preparation.

#### 2.6.2 LC-MS/MS Data Acquisition

Peptide samples were analyzed by liquid chromatography-tandem mass spectrometry (LC-MS/MS) using a Thermo Scientific Vanquish UHPLC system coupled to a Thermo Scientific Q Exactive mass spectrometer. Peptides were separated on a HALO ES-C18 column (160 Å, 2.7 µm, 1.0 × 250 mm; Advanced Materials Technology, Part No. 92121-902) at a flow rate of 50 µL/minute using a binary gradient with mobile phase A (0.1% formic acid in water) and mobile phase B (acetonitrile containing 0.1% formic acid). The gradient was as follows: 2% B from 0 to 5 min; linear ramp from 2% to 35% B over 5–50 min; 35% to 50% B from 50–53 min; 50% to 90% B from 53–55 min; held at 90% B from 55–57 min; returned to 2% B from 57–67 minutes for column re-equilibration; total run time was 70 min.

The Q Exactive was operated in positive ion mode with data-dependent acquisition (DDA). Full MS survey scans were acquired over a mass range of 200–2000 *m/z* at a resolution of 35,000 (at 200 *m/z*), with an automatic gain control (AGC) target of 5 × 10^5^ and a maximum injection time of 100 ms. The top 10 most abundant precursor ions (TopN = 10) were selected for fragmentation with a loop count of 10. MS² spectra were acquired at a resolution of 17,500, with an AGC target of 1 × 10^5^ and a maximum injection time of 50 ms. Precursor ions were isolated using a 4.0 *m/z* isolation window and fragmented by higher-energy collisional dissociation (HCD) at a normalized collision energy (NCE) of 30. The minimum AGC threshold for triggering MS² was set to 8.0 × 10^3^ with an intensity threshold of 1.6 × 10^5^. Peptide match was set to preferred, isotope exclusion was enabled, and dynamic exclusion was set to 10 s. Default charge state was set to 2+.

#### 2.6.2 Proteomics Data Analysis

Raw MS data were processed using MaxQuant (version 2.7.5.0). Peak lists were searched against the UniProt human reference proteome (UP000005640, Homo sapiens, taxonomy ID 9606) supplemented with the MaxQuant contaminants database. Trypsin was specified as the protease with up to two missed cleavages permitted. Carbamidomethylation of cysteine was set as a fixed modification. Variable modifications included oxidation of methionine and proline, and trioxidation of cysteine. Minimum peptide length was set to 4 amino acids and maximum peptide mass to 4,600 Da. Peptide-spectrum matches and protein identifications were filtered at a false discovery rate (FDR) of 1% at both the peptide and protein levels. Label-free quantification (LFQ) was performed within MaxQuant using the LFQ algorithm with a minimum ratio count of 1. LFQ intensities, protein groups, peptide groups, and gene-level identifications were exported from the MaxQuant output for downstream analysis. LFQ intensities were used for protein fold change calculations with non-transfected cells serving as the control group against which the transfected groups were compared.

### 2.7 Phenotypic characterization of primary human macrophages via flow cytometry analysis

Human peripheral monocytes were differentiated to macrophages (monocyte-derived macrophages, MDMs) by seeding at 10^5^ cells/cm^2^ in non-treated polystyrene 6 well-plates and cultured in RPMI 1640 media supplemented with 10% FBS, 1% pen/strep, and 25 ng/mL macrophage colony stimulating factor (M-CSF) (PeproTech) (complete macrophage media) for one week.^39^ MSCs were transfected in transwell inserts with a pore size of 0.4 μm and culture area of 4.1 cm^2^ and placed over the MDMs for co-culture. MDMs were co-cultured or polarized for 48 hours, then 1x brefeldin A was added to the media overnight. The cells were lifted by incubating in PBS supplemented with 5 mM EDTA on ice for 30 minutes, scraped gently, then washed thoroughly to remove remaining cells. The cells were washed in cell staining buffer (BioLegends) then stained with the LIVE/DEAD Fixable Yellow Dead Cell Stain Kit (Invitrogen) according to the manufactures protocol. The cells were washed then incubated with Human TruStain FcX (BioLegend) for 15 minutes at 4°C to block Fc receptors. Cell surface stains CD86 (B7-2) monoclonal antibody (IT2.2), Super Bright 436 (Invitrogen), CD206 (MMR) monoclonal antibody (19.2), Alexa Fluor™ 488 (Invitrogen), and CD163 monoclonal antibody (MAC 2-158), NovaFluor™ Red 725 (eBioscience) were then added at the optimized dilutions (Table S3) and incubated for 30 minutes at 4°C. The cells were fixed and permeabilized using the Cytofix/Cytoperm kit (BD). All data was collected on a cytoflex flow cytometer (Beckman Coulter, USA) and analyzed using FlowJo v10.10.0. Compensation was performed on single-stain controls using cells treated with lipopolysaccharide (LPS) (100 ng/mL) (eBioscience) and interferon-γ (100 ng/mL) (PeproTech) for CD86 or interleukin 4 (IL-4) (25 ng/mL) (PeproTech) and IL-13 (25 ng/μL) (PeproTech) for CD206 and CD163. Gating was performed on fluorescent minus one (FMO) controls and gating strategies can be visualized in Fig. S5.

### 2.7. Statistics

Statistical analyses were performed using GraphPad Prism (version 11.0.0) software. All data were tested for normality using the Shapiro–Wilk test. For normal data with three or more groups, a one-way analysis of variance (ANOVA) was used with a Brown–Forsythe test and a Tukey post-test. Non-normal data were analyzed using the Kruskal–Wallis test with multiple comparisons via Dunn’s test. Multi-day experiments were analyzed by two-way ANOVA with Tukey post-test. Numerical and graphical results were displayed as mean ± standard deviation (SD) for Sanger sequencing data, transfection efficiencies, flow cytometry median fluorescent intensities, and alamarBlue data. Log_2_ fold changes (log_2_FC) are presented as estimated fold change ± standard error (SE). Significance was accepted at *p* < 0.05. Sample size (*n*) is indicated within the corresponding figure legends. Each datapoint represents an individual sample from a single experiment.

## 3. Results

### 3.1 GFP mRNA delivery reveals distinct transfection profiles in MSCs across non-viral delivery systems

To establish the delivery capabilities of selected non-viral CRISPR delivery systems, we first transfected MSCs with GFP mRNA using CPPs, lipids, polymers, or electroporation and assessed transfection efficiency by monitoring GFP-fluorescence. **(Fig. 1A)**. Electroporation protocols were optimized to maximize the overall transfected cell yield (percent transfected x percent viable) (Fig. S6). CPPs and lipids yielded much higher transfection percentages than polymers and electroporation, transfecting 85.5 ± 2.18 and 93.2 ± 0.35 percent of MSCs respectively (**Fig. 1B**). Although both CPPs and lipids gave high transfection percentages, the median fluorescent intensity following lipid-based transfection was 19.44 ± 2.92-fold higher than CPPs and 414.15 ± 48.1 or 84.00 ± 7.94-fold higher than polymers and electroporation respectively **(Fig. 1C**). Conversely, CPP transfection minimized cytotoxicity with viability increasing from 84.41 ± 4.06 % on day 1 to 91.90 ± 3.55 % on day 3, compared to lipids which decreased viability from 70.03 ± 8.04 % to 63.44 ± 6.26 % from days 1 to 3 (**Fig. 1D**). Polymer and electroporation group increased from days 1 to 3 but remained significantly lower than the CPP group. These findings on the transfection percentages, MFI, and viability are also reflected in fluorescent images (**Fig. 1E).**

**Figure 1.**
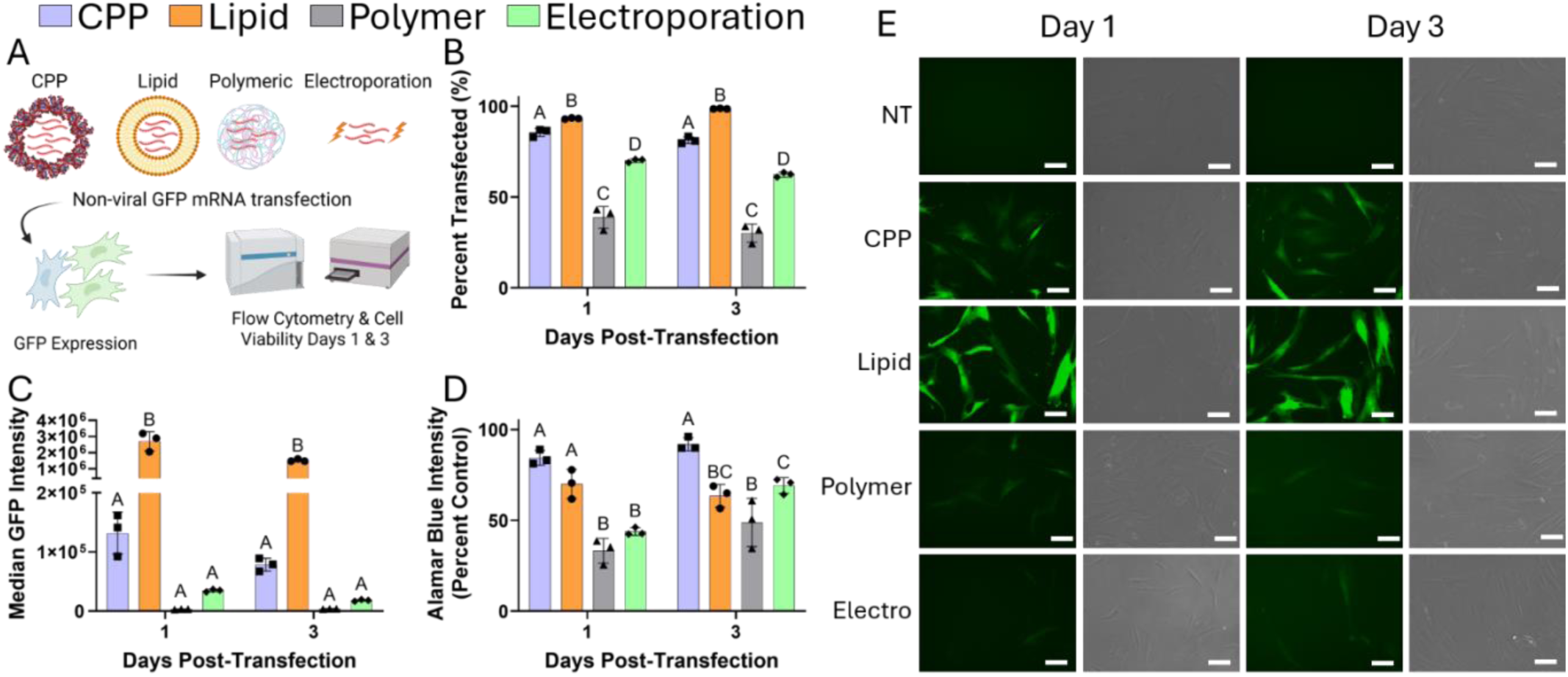
Non-viral transfection systems yield distinct transfection efficiencies and expression profiles. **A)** MSCs were transfected with GFP mRNA using the CPPs, lipids, polymers, or electroporation and characterized on days 1 and 3. Flow cytometry and alamarBlue were used to quantify **B)** the percentage of cells expressing GFP, **C)** the median fluorescent intensity of each population, and **D)** cell viability. Statistically similar groups at each timepoint are denoted with the same letters (two-way ANOVA, p < 0.05). **E)** Fluorescent images confirm the trends in transfection percentage, GFP brightness, and cell viability (NT = non-transfected). Scale bar = 100 μm.

### 3.2 Editing efficiency is determined by the choice of non-viral delivery system and CRISPR molecular format

To directly compare the CRISPR gene editing efficiencies across CRISPR molecular formats and delivery systems, we targeted the *AAVS1* safe-harbor locus using co-delivered Cas9 mRNA and gRNA (mRNA) or pre-complexed Cas9 and gRNA (RNP) (**Fig. 2A**). Cells were cultured for one week and analyzed by Sanger sequencing at the cleavage site. Our results indicate that lipid-based mRNA delivery systems yielded the highest overall editing efficiency with an 83.6 ± 0.4 % indel rate (**Fig. 2B**). However, this rate was not maintained with lipid-based RNP delivery with only 21.2 ± 1.6 % indels (**Fig. 2C**). Alternatively, CPPs achieved gene editing in both mRNA and RNP formats with 36.9 ± 1.9 and 49.2 ± 4.7 % indels respectively. In contrast, polymeric nanoparticles and electroporation featured lower editing rates overall (**Fig. 2B-C**). The elevated editing rates following CPP and lipid transfection directly correlate to the transfection rates and MFIs observed in **Fig. 1**. Additionally, editing via CPPs, polymers, and electroporation resulted primarily in precise 1 bp deletions, while lipid-based mRNA and RNP delivery yielded 15.29 ± 2.92 and 13.52 ± 0.35 % of larger indels respectively. These findings suggest that delivery system selection critically impacts genomic stability, highlighting inherent trade-offs between editing efficiency and genomic stability (**Fig. 2B-C**).

**Figure 2.**
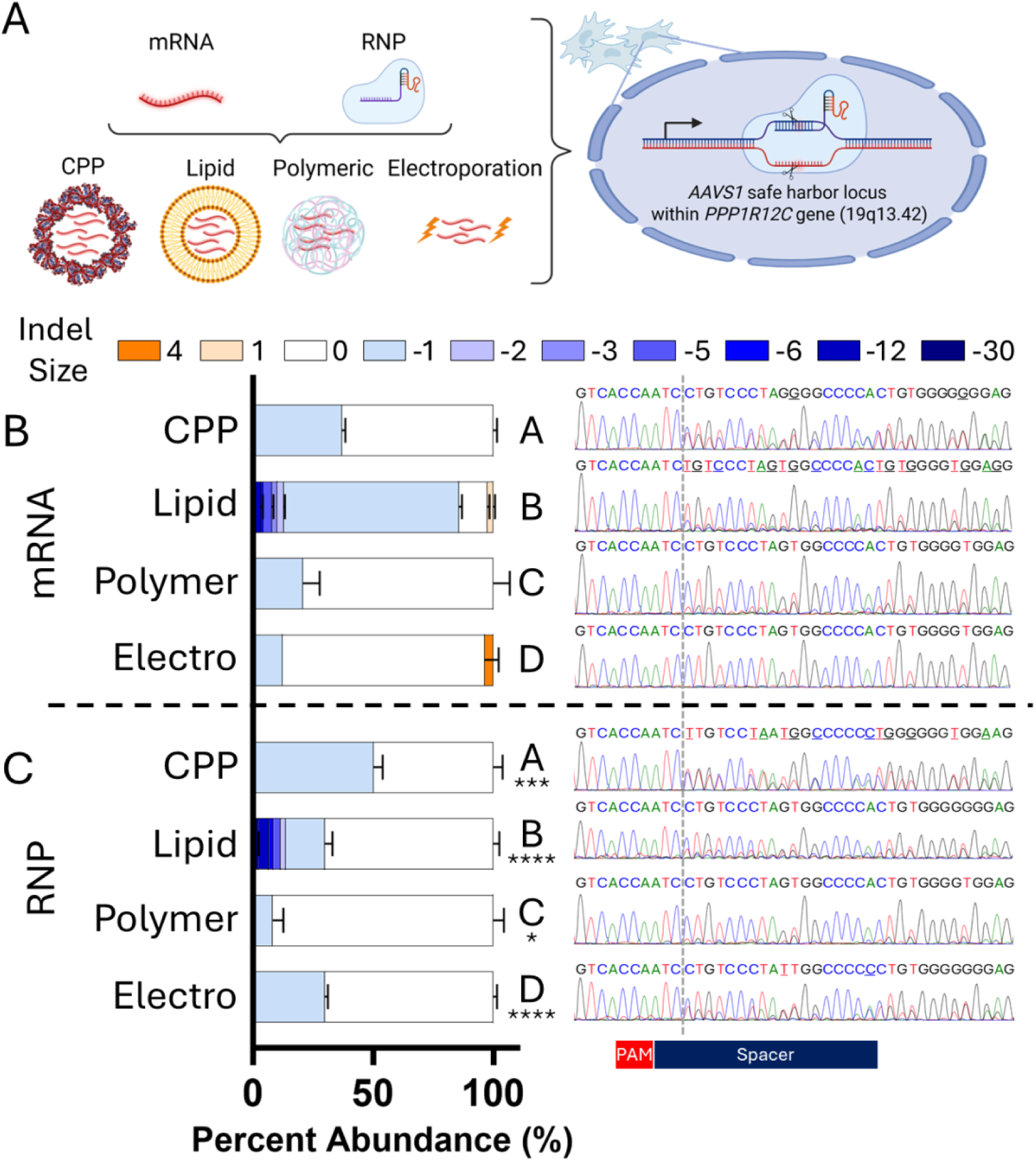
Editing efficiencies across CRISPR modalities and delivery systems. **A)** Schematic of experimental design. CPPs, lipids, polymers, or electroporation were used to knock-out the *AAVS1* safe harbor locus in MSCs by delivering CRISPR mRNAs or RNPs. Knock-out efficiencies were determined by Sanger sequencing for **B)** mRNA and **C)** RNP delivery and results are plotted as the relative abundance of each insertion/deletion (indel) mutation size (n=3). Groups with different overall editing efficiencies (p<0.05) do not share a letter. RNP groups that are significantly different from the corresponding mRNA group are also labelled * p<0.05, *** p<0.001, **** p<0.0001. Representative sanger sequencing traces are included for each group with the primary nucleotide signal labelled. Primary nucleotides that differ from the wild-type sequence are underlined.

### 3.3. RNA sequencing and differential expression analysis reveals distinct transcriptional dysregulation for each non-viral transfection system

Since high amounts of cell stress can interfere with double-stranded break (DSB) repair and influence indel mutation rates, we analyzed the transcriptomic profile of transfected cells to gain more insight into the potential dysregulation induced by each system.^40,41^ To better frame the impact of transfection on stem cell health, non-transfected MSCs were exposed to etoposide to serve as a senescent control. Dimensionality reduction and clustering analyses of RNA-seq counts were used to assess global transcriptomic differences between treatment groups. Following UMAP and tSNE, CPP-transfected and electroporated MSCs consistently clustered with non-transfected controls, whereas polymeric and lipid-based systems were separated (**Fig. 3A-B**). Additionally, k-means clustering after PCA demonstrates that the CPP-transfected group clustered most closely with non-transfected MSCs (**Fig. 3C**). Jensen-Shannon distance analysis corroborated these findings, with lipid-based systems exhibiting the greatest transcriptomic divergence from non-transfected controls (**Fig. 3D**). Together, these results consistently indicate that CPP and electroporation transfections cause the least transcriptomic disruption of the methods tested.

**Figure 3.**
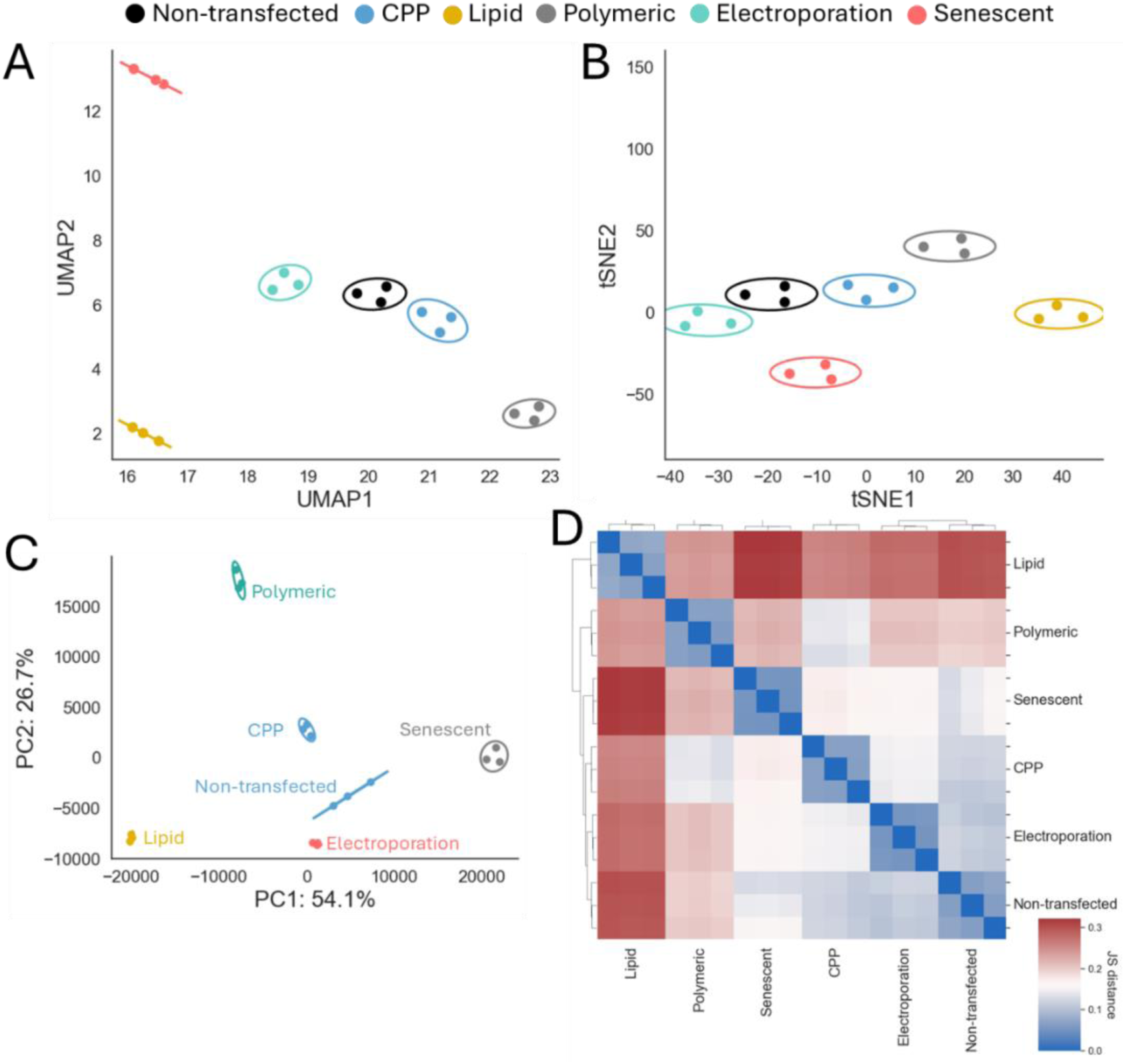
Global analysis of MSC transcriptomic profiles following non-viral transfection. Scatter plots showing a 2-dimensional **A)** UMAP and **B)** tSNE projections with confidence ellipses (2 standard deviations) overlaid for each cluster. **C)** Scatter plot of the first two principal components are colored by k-means clustering classification (k = 5) and overlaid with confidence ellipses (2 standard deviations). **D)** Heat map representing the Jensen-Shannon distance between each sample. Hierarchical clustering based on these distances is provided on the x and y axis.

To better understand the individual genes driving this transcriptomic dysregulation, we performed differential expression analysis relative to non-transfected controls for each transfection system (**Fig. 4**). CPP, lipid, and polymer systems upregulated overlapping pathways, with lipids causing the greatest dysregulation and CPPs consistently causing the least.

**Figure 4.**
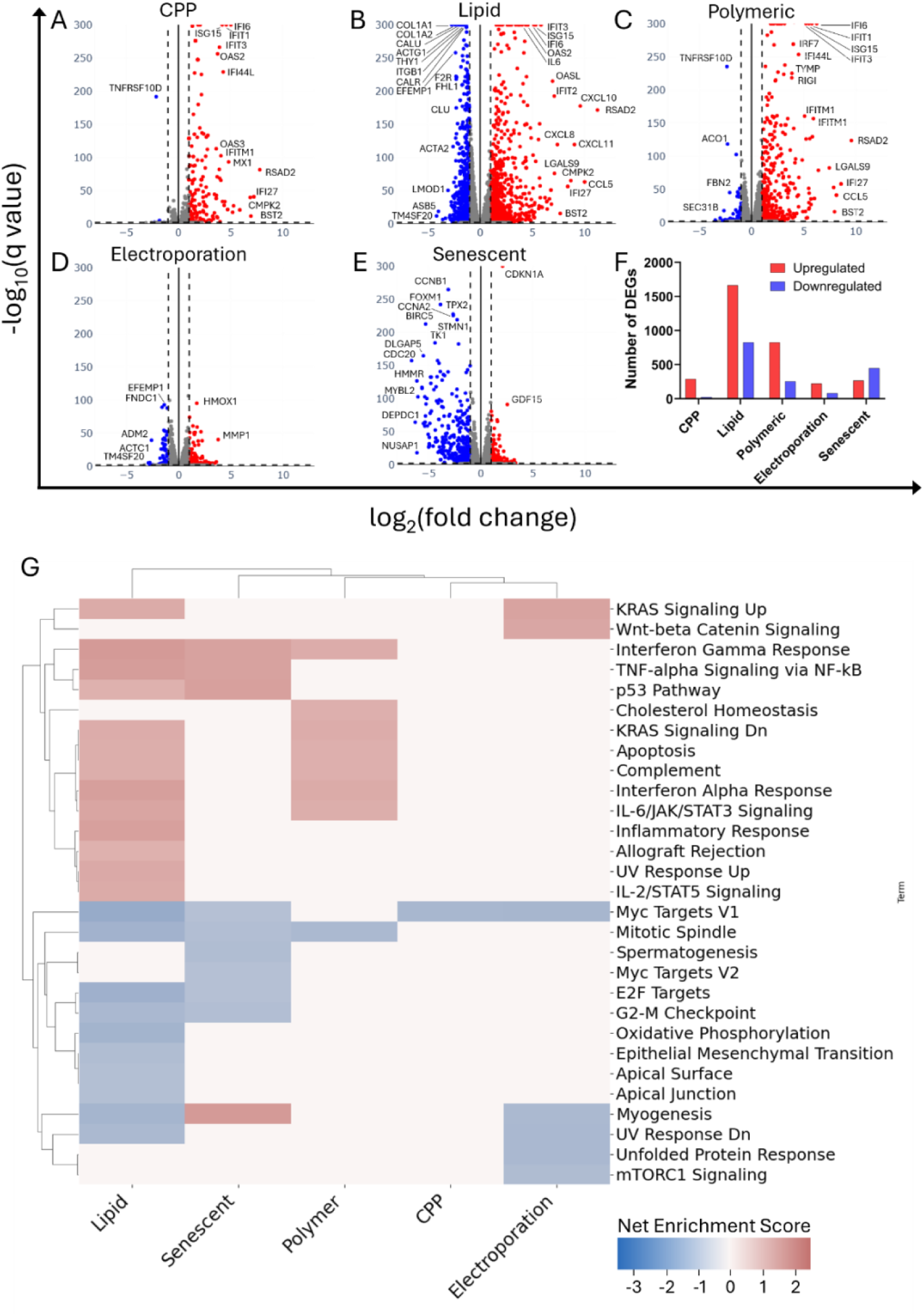
Non-viral transfection systems induce distinct transcriptomic dysregulation profiles. Volcano plots showing significant (q < 0.05, −1 > log_2_FC > 1) DEGs following transfection with **A)** CPPs, **B)** lipids, **C)** polymers, or **D)** electroporation compared to non-transfected controls. **E)** Volcano plots showing significant (q <0 .05, −1 > log_2_FC > 1) DEGs following treatment with etoposide to induce senescence. For all volcano plots, significantly upregulated genes (q < 0.05, log_2_ fold change > 1) are plotted in red, significantly downregulated genes (p<0.05, log_2_ fold change > 1) are plotted in blue, and non-significantly effected genes (q < 0.05, −1 < log_2_ fold change < 1) are plotted in gray. **F)** Bar plot of significant upregulated and downregulated (q < 0.05) DEG counts for each treatment based on differential expression analysis on n = 3 samples each. **G)** Cluster map of all gene ontology terms that were correlated (FDR < 0.25) to differential expression results for each treatment. Shade represents the net enrichment score (NES) resulting from gene set enrichment analysis (GSEA).

Differential expression signatures, specifically *TLR2* and *TLR4* upregulation **(Fig. 4)**, suggest that nanoparticles were first detected by toll-like receptors on the cell surface, initiating *MyD88* and Trif (*TICAM1)* signaling pathways which were further amplified by *TLR3* signaling from within the endosome from all nanoparticle strategies, initiating an NF-κB, and AP-1 signaling cascade. These pathways both independently and synergistically activate a range of inflammatory chemokines including IL-1β, TNF-α, IL-6, and CXCL10.^42–44^ Once within the cell, the nanoparticle reagents damaged the mitochondrial membrane and endoplasmic reticulum, causing the release of dsDNA and mitochondrial RNAs (mt-RNAs) into the cytoplasm. These ectopic nucleic acids were sensed by *RIGI* (RIG-I), *MDA5*, and cGAS-STING which subsequently signal through mitochondrial antiviral-signaling (MAVS) pathways to activate *IRF3* and *IRF7* transcription factors.^45,46^ IRF3 and IRF7 then enter the nucleus and activate the expression of anti-viral signaling pathways, primarily through type I interferons interferon-β (IFN-β) and interferon-α (IFN-α).^47,48^ Concurrently, ectopic dsDNA released from cellular damage activated the AIM2 inflammasome which increases caspase I activity and activates IL-1β into its functional isoform in a cell stress-dependent manner.^49^ This IL-1β activation is especially impactful for lipid transfections since *IL1B* was upregulated 84.68 ± 35.83-fold compared to 1.44 ± 0.61, 1.91 ± 0.81, and 1.46 ± 0.62-fold for CPPs, polymers, and electroporation respectively.

In addition to secreting these inflammatory factors, transfected MSCs upregulated the receptors for interferon-β, IL-1β, and TNF-α which created an inflammatory feed-forward loop, that was most prevalent with lipid transfected cells. Ligand binding to these surface receptors continuously stimulates both canonical and non-canonical NF-κB pathways, IRF1, IRF7, interferon-λ, and a range of interferon stimulated genes (ISGs). Importantly, IRF1 is also known to drive its own autocrine loop characterized by sustained expression of ISGs and inflammatory chemokines.^50,51^ Among these ISGs, the interferon-induced proteins with tetratricopeptide repeats (IFIT) genes were highly upregulated due to their abilities to bind exogenous or ectopic RNAs and amplify interferon signaling. For example, *IFIT3* which binds RNAs with short untranslated regions (UTRs) and amplifies IRF3 signaling^52^, was upregulated 15.16 ± 1.18, 56.89 ± 4.41, 37.07 ± 2.87, and 1.32 ± 0.10-fold in CPPs, lipids, polymers, and electroporation respectively (**Fig. 4A–C**). 2’-5’-oligoadenylate synthetase (OAS) proteins then sense exogenous RNAs and activate RNase L to degrade them. OAS3, a primary activator of RNase L and the dominant OAS isoform responsible for antiviral RNA degradation,^53,54^ was upregulated 12.34 ± 1.33, 23.08 ± 2.49, and 16.61 ± 1.79-fold for CPPs, lipids, and polymers respectively. Additionally, *OASL* which enhances RIGI signaling was upregulated 15.32 ± 2.34, 121.55 ± 18.56, and 60.63 ± 9.26-fold respectively.^55^ Many other ISGs followed these same trends including *MX1*, *RSAD2*, and *ISG15* which inhibit transcription, terminate translation, and tag proteins to reduce their activity respectively.^56–58^

Inflammatory signaling is also known to license MSCs for immunomodulation to remodel their environment and counteract inflammation.^59^ On the transcriptomic level, lipid-transfected MSCs appear to be licensed for immunomodulation with elevated expression of *IDO1*, *PGE2*, *TNFAIP6* (TSG-6), and *CD274* (PD-L1).^60,61^ Additionally, high *IRF1* and *NLRC5* expression stimulates the production of MHC-I molecules HLA-A and HLA-B which inhibit continued TLR activation and protect MSCs from natural killer (NK) cell targeting.^55–57^ Yet, secreted inflammatory signals still dominate these signals, specifically for lipid transfected cells. *CXCL10*, a pleiotropic CXC chemokine that recruits T cells, macrophages, and NK cells via CXCR3, was upregulated 18.69 ± 4.49, 764.81 ± 183.88, 84.48 ± 20.31, and 1.00 ± 0.24-fold for CPPs, lipids, polymers, and electroporation respectively.^62^ Similarly, *CCL5*, a chemokine constitutively secreted by MSCs that recruits macrophages and T cells via CCR5,^63–65^ was upregulated 57.71 ± 23.29, 994.49 ± 401.24, 73.31 ± 110.27, and 4.03 ± 1.63-fold respectively. Other potent inflammatory genes including *CXCL8, CXCL9*, *CXCL11*, *TNFSF10* (TRAIL), *LIF,* and *TNFSF13B* (BAFF) were also upregulated in similar patterns. Furthermore, both *CSF2* (GM-CSF) and *CSF3* (G-CSF) which polarize macrophages to inflammatory and pro-regenerative phenotypes respectively are highly upregulated following lipid transfection.^66,67^

Electroporation minimally affected antiviral ISG and NF-κB pathways due to a lack of TLR signaling and mitochondrial disruption. However, electroporation uniquely dysregulated matrix metalloprotease (MMP) genes. Electroporation upregulated MMP1 and MMP3 by 13.63 ± 2.57 and 4.12 ± 1.62-fold respectively, comparable only to lipid-based systems which upregulated them 7.44 ± 1.40 and 6.03 ± 2.36-fold respectively, while CPP and polymer systems did not significantly affect MMP expression. These MMPs were part of a broader transcriptional disruption that moderately stimulated Wnt/β-Catenin pathways while negatively enhancing mTORC1 and RAS signaling (**Fig. 4G**).

Finally, MSC senescence was induced through treatment with etoposide, a topoisomerase II inhibitor.^68,69^ Etoposide treatment primarily downregulated gene expression, with 449 significantly downregulated genes (p_adj_ <0.05, log_2_FC < −1) primarily involved in cell cycle and mitotic progression (**Fig. 4E)**. In comparison, liposomes significantly downregulated 827 genes compared to 24, 255, and 82 for CPPs, polymers, and electroporation respectively (**Fig. 4F**). Among the downregulated genes from lipid-based nanoparticles, 145 were shared with senescent MSCs, with the most downregulated genes being *KIF20A*, *ANGPTL, CDC20, CDK15 and CDCA3.* Alternatively, CPPs, polymers, and electroporation shared 1, 17, and 1 downregulated gene in common with senescent MSCs. These relationships are also reflected in gene set enrichment analysis (GSEA) with lipids and senescent controls sharing in the negative enrichment of Myc targets, E2F targets, G2/M checkpoint, and mitotic spindle (**Fig. 4G**).

GSEA also demonstrated that lipids and polymers enhanced type I interferon signaling, apoptosis, the complement system, and IL-6/JAK/STAT pathways, which can be attributed to their increased activation of TLRs and RIG-I pathways (**Fig. 4G**). Additionally, lipids specifically enriched TNF-α signaling and inflammatory response confirming that it induced the greatest overall inflammatory response. Transfection also disrupted the MSC transcriptome beyond inflammatory signaling. All transfected groups except for polymeric downregulated the MYC targets V1 gene set which can be attributed to the downregulation of translation factors *EEF1B2*, *PABPC1*, and *PABPC4* and ribosome biogenesis proteins *RACK1*, *ODC1*, and *IMPDH2*.^70^ The downregulation of these genes is inversely correlated to the fluorescent intensities of transfected eGFP mRNA indicating the cell’s response to an overabundance of the transgene and resulting proteins. Additionally, lipid and polymer groups negatively enriched the mitotic spindle gene set through the downregulation of *ANLN*, *CENPF*, *KIF23*, *SMC4*, *TOP2A*, *TPX2*, which have also been identified as key stemness-associated genes.^71,72^ Lipids also shared the negative enrichment of G2-M checkpoint genes with the senescent control indicated reduced cell division and proliferation which is an essential stem cell property, a requirement for CRISPR knock-in strategies, and an important step in achieving functional edited populations.^34,73,74^

### 3.4 DDA proteomics confirms RNA-seq trends

While RNA sequencing reveals distinct inflammatory signatures, many of these pathways rely on phosphorylation or post-translational modifications, necessitating protein-level confirmation (**Fig. 5A**). Bottom-up DDA proteomics confirmed the enrichment of ISGs including IFIT1, IFIT2, IFIT3, and IFITM1 in both lipid and polymer groups but not CPPs or electroporation, consistent with gene expression levels (**Fig. 5B**). Several other interferon-related proteins were specifically enriched following lipid transfection including STAT1, OASL, RIG-I, RSAD2, B2M, and CCL5 confirming the robust and sustained inflammatory response (**Fig. 5B**). Notably, the enrichment of STAT1 confirms a prolonged type 1 interferon response and subsequent activation of JAK/STAT pathways which drives downstream ISG production, licenses MSC immunomodulation, and impairs differentiation.^75–77^

**Fig. 5.**
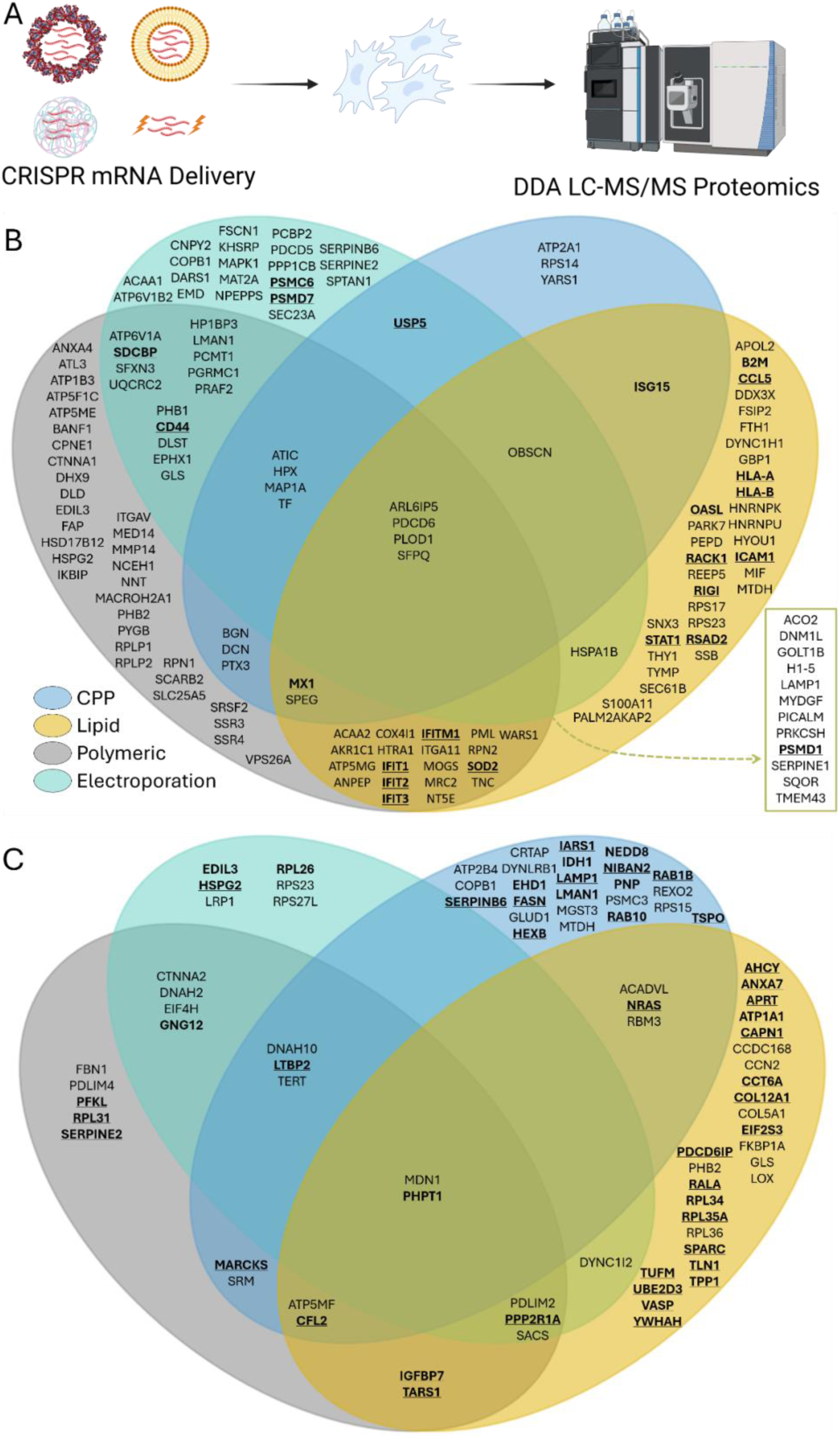
Proteomic analysis of transfected cells. **A)** MSCs transfected with Cas9 mRNA and *AAVS1* gRNA were analyzed via bottom-up DDA proteomics. **B)** Proteins that were enriched in each sample but were not detected in each sample were plotted in a venn diagram to visualize the overlap between samples. Proteins are listed by their gene names and proteins mentioned in the text are in bold. **C).** Proteins that were identified in non-transfected controls but not detected in each sample are compared on a venn diagram. Proteins that contribute to the extracellular exosome (GO:0070062) or extracellular vesicle (GO:1903561) gene sets in the cellular components gene ontology are listed in bold.

Lipids and CPPs, but not polymers, enriched ISG15, an interferon-stimulated ubiquitin-like modifier that tags antiviral proteins through a process called ISGylation that enhances autophagy pathways.^58^ Increased ISG15 correlates to the copy number of transfected mRNAs suggesting that CPP and lipid transfected cells are responding to the abundance of non-self-proteins. This enrichment is also accompanied with two unique sets of proteins involved in protein transport, MVB formation, lysosomes, and EV secretion that were not detected in CPPs and lipids respectively (**Fig. 5C**).^78^ The distinction between these protein sets indicates different stages of cell recovery. CPPs had more abundant USP5 which inhibits ISG15 and de-ISGylates proteins suggesting the end of autophagy pathways.^79^ Alternatively, lipid transfection enhanced the complement system autophagy pathway, upregulating *C3* 5.26 ± 1.87-fold and significantly upregulating *MAP1LC3B* and *ATG16L1* gene expression.^80^ The combination of ISG15 and complement pathways likely amplified autophagy leading to a prolonged response. Importantly, autophagy mechanisms are known to interfere with EV secretion pointing to potential disruption of important MSC immunomodulatory pathways.^81^

Alternatively, polymeric nanoparticles and electroporation shared enriched proteins associated with EV secretion. For example, both groups demonstrated increased levels of syntenin (*SDCBP*), a central regulator of exosome biogenesis and secretion^82,83^ This is accompanied by higher abundance of CD44 which is a well-characterized exosome surface marker and cargo.^84^ Additionally, electroporated cells featured highly abundant proteasomal subunits PSMC6, PSMD7, and PSMD1 indicating upregulated proteolysis in response to damaged protein which can result in the secretion of damaged proteins through EVs.^85^ USP5 is also enriched following electroporation and likely contributes to this protein remodeling process.^79^ Alternatively, polymer transfection enriched mitochondrial and endoplasmic reticulum proteins indicating that the surviving cells are replacing damaged proteins lost during mitochondrial/ER damage (Fig. 5B). EV secretion also might be responsible for the lack of IGFBP7 detection after polymer transfection. While IGFBP7 protein was not detected in lipid or polymer groups, its gene expression was significantly upregulated 1.66 ± 0.05-fold from a very high basal expression with polymers and significantly downregulated 0.67 ± 0.02-fold with lipids. This drastic change indicates that polymer transfected groups may be secreting anti-inflammatory cues in response to transfection^86^.

To further visualize the underlying molecular mechanisms, we mapped the impact of each delivery system on key inflammatory genes (**Fig. 6**).

**Figure 6.**
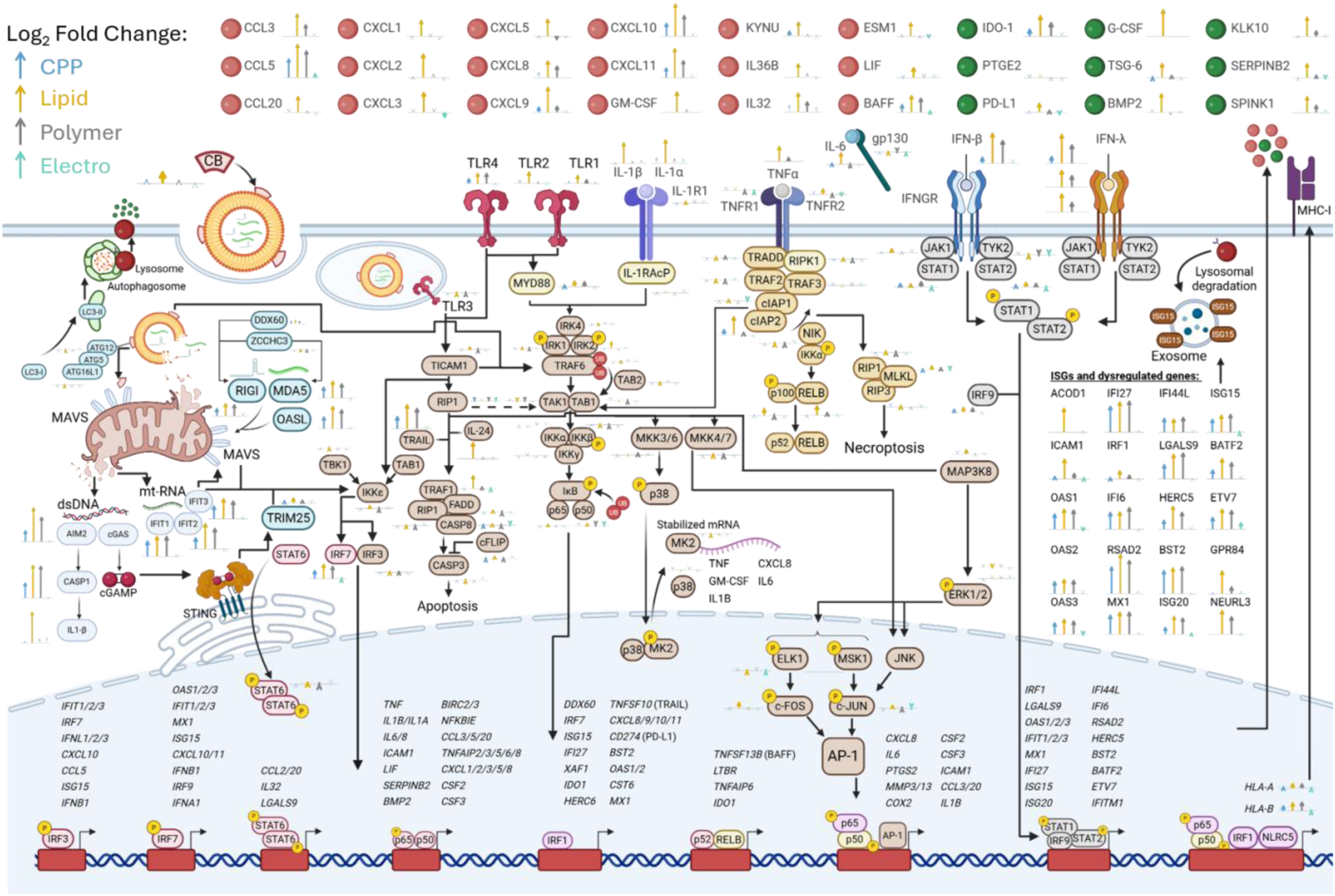
Proposed mechanisms of transcriptomic dysregulation following non-viral transfection. The log_2_ fold change in gene expression for each molecule after transfection with each system are represented by arrows with significance (q <0.05) indicated by the thickness of the line (thick lines indicate significance).

### 3.5 Non-viral CRISPR delivery disrupts the immunomodulatory capacity of transfected cells

Transcriptomic and proteomic evidence of altered EV biogenesis pathways suggests that transfection-induced stress may alter the immunomodulatory capacity of MSCs. MSCs are inherently immunomodulatory, and when primed with inflammatory stimuli they acquire enhanced immunosuppressive capacity.^87^ However, excessive or dysregulated stimulation can shift MSCs toward a pro-inflammatory phenotype characterized by interferon-stimulated gene upregulation, chemokine secretion, and impaired paracrine immunomodulation.^88^ Given the conflicting immunomodulatory profiles identified in RNA-seq and proteomic analysis, we sought to determine the impact of CRISPR delivery on the innate immune system using a primary MDM co-culture system. MSCs were transfected with Cas9 mRNA and gRNA targeting the *AAVS1* safe-harbor locus using CPPs, lipids, polymers, or electroporation and subsequently co-cultured with MDMs for 48 hours (**Fig. 7A**). Immunostaining and flow cytometry were performed to characterize MDM expression of CD86, CD206, and CD163 with non-treated MDMs and MDMs co-cultured with non-transfected MSCs serving as controls.

**Figure 7.**
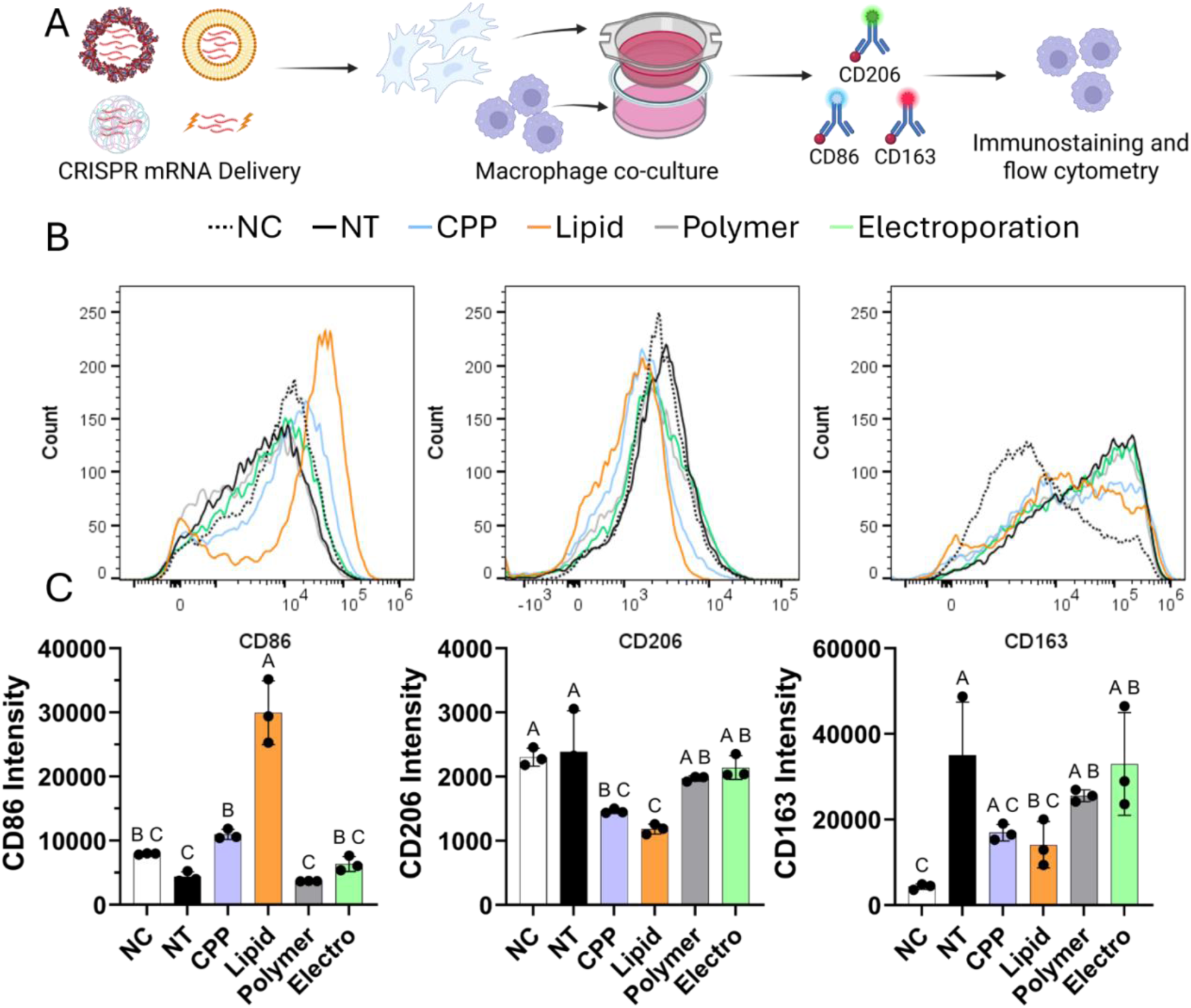
Effects of CRISPR delivery method on MSC immunomodulatory properties. **A)** MSCs were transfected with Cas9 mRNA and *AAVS1* gRNA using CPPs, lipids, polymers, or electroporation then co-cultured with primary human M0 MDMs for 48 hours. The resulting MDM phenotypes were assessed by immunostaining and flow cytometry. **B)** Representative histograms of surface marker expression shown as a single concatenated file of n = 3 samples, down sampled for consistency. NC represents non-treated M0 MDMs and NT represents M0 MDMs co-cultured with non-transfected MSCs. **C)** The corresponding median fluorescent intensity ± standard deviation for each condition (n = 3). Samples that are not statistically significantly different (p < 0.05) as determined by a one-way ANOVA are denoted by the same letter.

Surface marker analysis on co-cultured MDMs revealed distinct immunomodulatory effects from each transfection system. CD86 is typically upregulated in pro-inflammatory macrophages and is involved in antigen presentation.^89^ Co-culture with non-transfected MSCs non-significantly reduced CD86 expression, whereas lipid transfection significantly upregulated CD86 3.77 ± 0.51-fold (**Fig. 7B-C**). Although co-culture with CPP-transfected MSCs did not significantly increase CD86 expression compared to non-treated MDMs, it resulted in significantly higher CD86 expression compared to co-culture with non-transfected MSCs. CD206, which is most upregulated by IL4-stimulated macrophages and is associated with tissue repair, was significantly reduced 0.63±0.02 and 0.51±0.03-fold during co-culture with CPP and lipid-transfected MSCs respectively (**Fig. 7B-C**).^33,89^ Similarly, CD163, which is upregulated by IL10-stimulated macrophages, was significantly upregulated 15.20±4.40-fold during co-culture with non-transfected MSCs but was not significantly increased during co-culture with CPP or lipid transfected MSCs (**Fig. 7B-C**).^33^ Polymer and electroporation transfection systems did not significantly alter the expression of surface markers in co-cultured MDMs. Taken together, this data indicates altered immunomodulatory capacity in MSCs transfected with CPP and lipid systems, with lipids exclusively upregulating the expression of the pro-inflammatory CD86 marker.

### 3.6. Decoupling delivery method from CRISPR molecular format reveals distinct transcriptomic effects in primary human MSCs

After analyzing the effects due to delivery method, we sought to evaluate the transcriptomic effects of CRISPR molecular format in MSCs. Our CPP system offers a unique platform for decoupling CRISPR molecular format from delivery system due to its versatility and efficient delivery of CRISPR mRNAs and RNPs **(Fig. 2).** Accordingly, we compared the transcriptomic profiles following CRISPR delivery in Cas9 mRNA versus RNP molecular formats (**Fig. 8A**). Compared to mRNA delivery, Cas9 RNP upregulated inflammatory chemokines, upregulating *CCL5* 3.52 ± 0.20-fold and *CXCL10* 2.57 ± 0.35-fold consistent with RIG-I activation from elevated gRNA concentrations (**Fig. 8B**).^90^ Consistent with GFP mRNA delivery, Cas9 mRNA and *AAVS1* gRNA only negatively enriched Myc Targets V1 (**Fig. 8C**). However, the changes in DEG expression with RNP delivery impacted GSEA results, with RNP transfection enriching KRAS Signaling down, Complement, IL-6/JAK/STAT3 Signaling, and inflammatory Response gene sets (**Fig. 8C**). These results demonstrate that RNP delivery increases inflammatory signaling in MSCs which could be attributed to the increase in RIGI pathways due to slightly higher gRNA concentrations.

**Figure 8.**
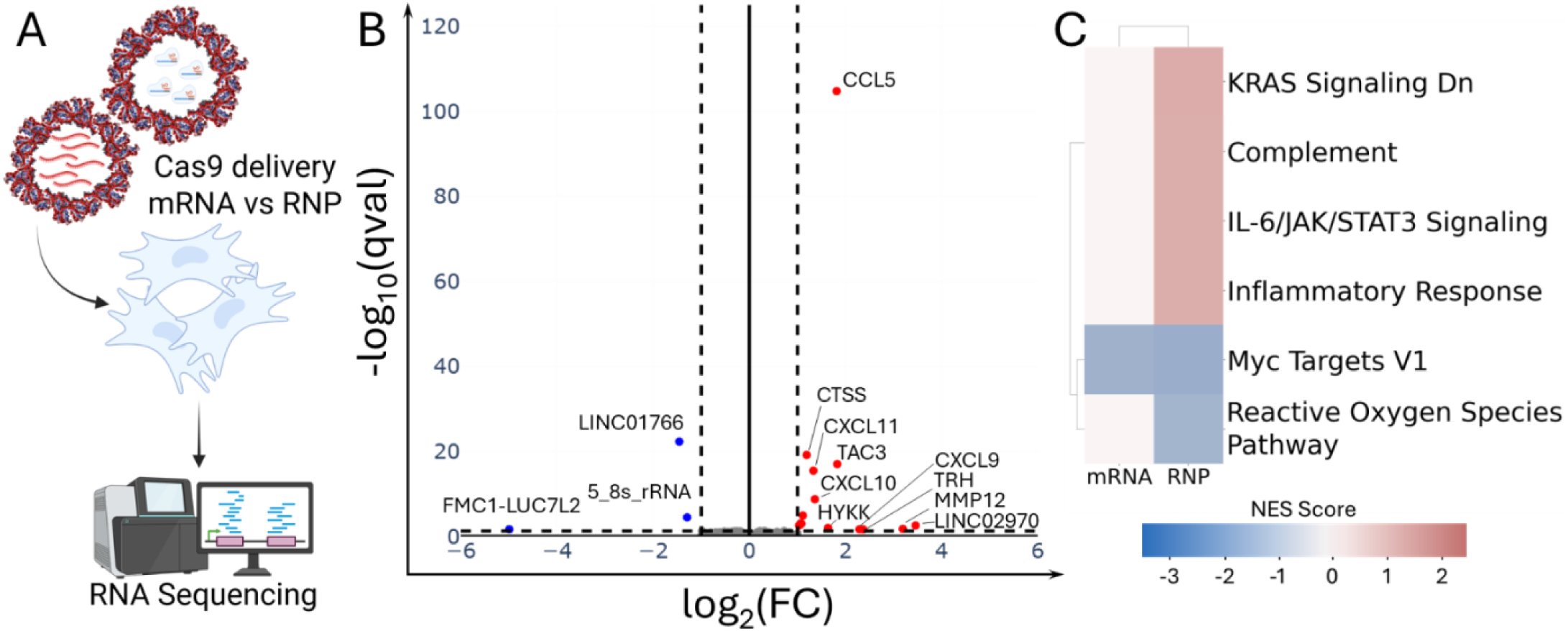
Transcriptomic analysis of CRISPR cargos. **A)** Schematic of experimental approach. MSCs were transfected with Cas9 mRNA or Cas9 RNP using CPPs before transcriptomic profiling via RNA-seq. **B)** Volcano plot representing the differential expression results for Cas9 RNP vs Cas9 mRNA delivery. Genes significantly upregulated in RNP (p<0.05, log_2_ fold change > 1) are plotted in red, significantly downregulated genes (p<0.05, log_2_ fold change > 1) are plotted in blue, and non-significantly affected genes (p<0.05, −1 < log_2_ fold change < 1) are plotted in gray. **C)** GSEA results for enriched MSigDB_Hallmark_2020 gene sets plotted as the net enrichment score (FDR<0.25).

## 4. Discussion

We demonstrate that the selection of non-viral delivery system and CRISPR molecular format significantly influences the safety and efficacy of gene editing in primary human MSCs. First, we established non-viral delivery with non-viral CPP, lipid, polymer, and electroporation systems using eGFP reporter mRNA. We show that while CPPs and lipids can deliver mRNA to the highest percentage of cells, lipids deliver much higher mRNA copy numbers while CPPs preserve cell viability. Alternatively, polymers and electroporation yielded lower values in all categories. Then, we compared these transfection profiles to CRISPR editing efficiencies by delivering CRISPR mRNA or RNP with gRNA targeting the *AAVS1* safe harbor locus to decouple editing rates from downstream gene regulatory networks.^91^ We show that editing efficiencies highly depend on mRNA copy numbers, with lipid-based mRNA delivery providing the highest editing rates followed by CPPs while polymers and electroporation resulted in low editing efficiencies. However, these trends were not consistent with RNP delivery where CPPs outperformed all other delivery systems.

We also show that the transfection system, rather than the observed editing rates, correlated with increased indel sizes, with both lipid mRNA and RNP delivery resulting in over 13% of indels larger than 1 bp. Kosicki *et al.* demonstrated that this increase in the size of small indels to directly correlates to the frequency of large deletions, translocations, and non-contiguous lesions.^92^ This genomic instability was also observed by Park *et al.*, who found that when high editing efficiencies (96%) were achieved, 41.2% of edits caused deletions larger than 50 bp, causing large structural variants^93^. Although detecting indels of this size requires advanced single-molecule real-time sequencing rather than standard Sanger sequencing, their higher prevalence in stem cells compared to cell lines makes them a significant concern for tumorigenesis and unpredictable adverse effects.^93^ Since cellular stress is known to impact NHEJ rates and genomic instability, we performed bulk RNA-seq on transfected cells to understand the underlying stress induce by each transfection system.^40,41^ Global analysis via UMAP, t-SNE, K-means clustering, and Jensen-Shannon distances indicates that CPPs and electroporation least perturbed the MSC transcriptome. Differential expression analysis further confirmed that CPPs and electroporation resulted in the fewest number of differentially expressed genes. We then showed that all nanoparticle-based systems induced cell stress primarily through IFN-β IL-1β, and TNF-α signaling causing a feed-forward inflammatory loop but lipid transfection consistently upregulated these pathways the most indicating the highest cell stress.

The balance between CRISPR editing rates and inflammatory signaling represents a significant concern for MSC therapies. We show that the compounding of TLR signaling, IL-1, TNF-α, and mitochondrial damage induce inflammatory cascades that outweigh the typical immunomodulatory licensing effects of MSCs. Although lipids and polymers express the highest levels of inflammatory markers, we show that lipid and CPP transfection have the most significant impact on MSC immunomodulation through the reduction of CD206 and CD163 expression. This observation can likely be attributed to the elevated ISG15 gene and protein levels in CPP and lipid transfected MSCs. ISG15 promotes the aggregation of multivesicular bodies (MVBs) with lysosomes, diverting their contents toward lysosomal degradation rather than extracellular secretion.^81^ This disruption of EV output and MSC-MDM cross-talk explains the reduced upregulation of pro-regenerative markers in co-cultured MDMs.^81^ Still, many upregulated inflammatory factors such as CXCL10 and CCL5 do not rely on exosomes for secretion, contributing to the strong upregulation of CD86 in MDMs co-cultured with lipid transfected MSCs. These factors risk local and systemic inflammation that can affect the safety and efficacy of an MSC therapy. For example, elevated CXCL10 levels have recently been linked to systemic inflammation and myocarditis following lipid-based nanoparticle COVID-19 vaccinations^62^. Other inflammatory factors including NF-κB, IL-1β, and TNF-α have also been demonstrated to reduce MSC proliferation rates, colony formation, and multipotency.^94–96^ These results highlight a distinct trade-off between editing rates, inflammatory signaling, and essential stem properties. Minimizing these effects must be considered since the paracrine suppression of local inflammation is essential for effective tissue regeneration and engraftment during MSC therapies.

RNA-seq analysis across transfection systems was performed by transfecting fully capped and pseudouridine (ψ) modified GFP mRNA. However, CRISPR RNAs, specifically gRNAs with 5’-triphosphate, robustly activate the RIG-I pathway.^24^ When comparing CRISPR RNP and mRNA transfection using our CPP system we demonstrate that RNP yields higher editing rates and higher activation of inflammatory markers which can be attributed to higher amounts of intracellular gRNAs. This increased inflammatory response helps to explain the loss of editing efficiency in lipid and polymer-transfected MSCs. Since lipid and polymeric groups induced the highest cell stress, their combination with RNP delivery likely exacerbates inflammatory signaling, increasing cell death and compromising editing rates.^24,97^ Alternatively, in the more cell-friendly transfection systems, RNP delivery was able to improve editing rates. Taken together these results demonstrate the importance of selecting the optimal transfection system for maximizing the therapeutic impact of a gene editing strategy.

This work elucidates many challenges in CRISPR-based MSC therapies that warrant further study. We show that at the delivered doses, CRISPR RNP delivery elicits greater inflammatory signaling than mRNA-based delivery. However, this effect can be largely attributed to slightly higher levels of gRNA transfected (5-fold) and the induction of *RIGI* signaling. Since Cas9 RNP is much smaller than Cas9 mRNA and excess gRNA is not needed for pre-complexed RNP, the same number of molecules can theoretically be delivered with less gRNA. Future work should investigate the trade-offs of editing rates and transcriptomic dysregulation at a range of RNP doses. Additionally, this work identifies multiple signaling molecules, namely IL-1β, IRF3, IRF7, STAT1, and IRF9 that are largely responsible for the inflammatory cascades after transfection. Future work will investigate the incorporation of small interfering RNAs (siRNAs) or small molecules to inhibit these signaling pathways during transfection to allow the uptake of CRISPR cargos and efficient editing with minimal cell stress. Finally, we characterized transcriptomic dysregulation two days post-transfection to elucidate more prolonged inflammatory pathways. We demonstrate that lipid transfection specifically led to autocrine loops and sustained activation of these pathways. However, the upregulation of specific genes such as *USP18*, *CFLAR*, and *SOCS3* might indicate that the cells are attempting to counteract these pathways.^98–100^ Additionally, the activation of USP5 in CPP groups could indicate the termination of ISG15 signaling and the return to normal EV secretion and immunomodulation in future time points.^79^ Future work will investigate the total duration of this inflammatory signaling to better understand the temporal dynamics of non-viral CRISPR delivery induced stress.

## 5. Conclusions

In conclusion, the selection of non-viral delivery system and CRISPR molecular format represents a critical determinant of gene editing outcomes in primary human MSCs. Transfection systems face fundamental trade-offs between editing efficiencies, cell viability, and cell stress that must be considered for effective MSC therapies. While lipid-based delivery achieves the highest editing rates, it induces the greatest genomic instability, transcriptional dysregulation, and inflammatory burden. This cell stress overwhelms the inherent immunomodulatory licensing capacity of MSCs, shifting them toward a pro-inflammatory phenotype characterized by secretion of mediators including CXCL10, with downstream consequences for proliferation, multipotency, and therapeutic utility. Conversely, CPP-based systems better preserve cell viability and transcriptomic integrity while achieving moderate editing efficiencies, whereas polymers and electroporation yielded low editing efficiencies. These findings collectively establish that editing efficiency alone is an insufficient metric for delivery system selection, and that genomic integrity, transcriptomic stability, and the preservation of stem cell immunomodulatory function must be considered as primary design criteria in the development of gene-edited MSC therapeutics.

## 6. Acknowledgements

This material is based upon work supported by the National Science Foundation under Engineering Research Initiation (ERI) Grant No. 2347637 to Tomas Gonzalez-Fernandez. Any opinions, findings, and conclusions or recommendations expressed in this material are those of the author(s) and do not necessarily reflect the views of the National Science Foundation. Additionally, Tomas Gonzalez-Fernandez would like to acknowledge the start-up funds provided by the Department of Bioengineering and the P.C. Rossin College of Engineering and Applied Science at Lehigh University, and the Career Development Award from the American Society of Gene and Cell Therapy. The content is solely the responsibility of the authors and does not necessarily represent the official views of the American Society of Gene and Cell Therapy. Josh Graham would like to acknowledge the support provided through the National Science Foundation Graduate Research Fellowship under Grant No 2234658. The authors gratefully acknowledge the use of the LCMS facility at the HEAL Center, College of Health, Lehigh University.

