## Supplemental Figures for "Integrative Multiomic Analysis Reveals How Non-Viral Delivery System Selection Shapes CRISPR Gene Editing Outcomes in Stem Cells"

**SUPPLEMENTAL DATA**

**Tables:**

Table S1: Thermocycler conditions for amplifying genomic DNA	1

Table S2: Gene Set definitions for GSEA	2

Table S3: Flow cytometry antibody information and dilutions	3

**Figures:**

Figure S1: Optimized electroporation protocols	4

Figure S2. PCR products for sanger sequencing	5

Figure S3: UMAP Dimensionality reduction after PCA	6

Figure S4: Optimized ‘k’ for k-means clustering analysis	7

Figure S5: Gating strategy used for flow cytometry	8

Figure S6: Transfection yields following non-viral transfection	9

**Table 1**. Thermocycler conditions for amplifying genomic DNA.

| Step | Temperature (˚C) | Time (s) | Cycles |
| --- | --- | --- | --- |
| Initial Denature | 98 | 60 | 1 |
| Denature | 98 | 15 | 30 |
| Anneal | 68 | 18 |  |
| Extension | 72 | 18 |  |
| Final Extension | 72 | 300 | 1 |

**Table S2.** Gene sets from "MSigDB_Hallmark_2020" identified by GSEA and corresponding definitions.

| Gene Set | Definitions |
| --- | --- |
| Wnt-beta Catenin Signaling | Wingless-related integration site (Wnt) |
| Interferon Gamma Response | N/A |
| p53 Pathway | N/A |
| TNF-alpha Signaling via NF-kB | Tumor necrosis factor alpha (TNF-α) / nuclear factor kappa B (NF-κB) |
| Inflammatory Response | N/A |
| KRAS Signaling Up | Kirsten rat sarcoma viral proto-oncogene (KRAS) |
| KRAS Signaling Down | N/A |
| Apoptosis | N/A |
| IL-6/JAK/STAT3 Signaling | Interleukin-6 (IL-6) / Janus kinase (JAK) / signal transducer and activator of transcription 3 (STAT3) |
| Complement | N/A |
| Allograft Rejection | N/A |
| Myogenesis | N/A |
| Epithelial Mesenchymal Transition | N/A |
| Unfolded Protein Response | N/A |
| mTORC1 Signaling | Mechanistic target of rapamycin complex 1 (mTORC1) |
| Myc Targets V1 | N/A |
| UV Response Dn | ultraviolet (UV) response, downregulated (Dn) |
| Spermatogenesis | N/A |
| Mitotic Spindle | N/A |
| Oxidative Phosphorylation | N/A |
| Apical Surface | N/A |
| Apical Junction | N/A |
| E2F Targets | E2 transcription factor (E2F) |
| G2-M Checkpoint | Gap 2-Mitosis (G2-M) |
| Reactive Oxygen Species Pathway | N/A |

**Table S3.** Flow cytometry antibody information and dilutions.

| Target | Fluorophore | Ex./Em. | Catalog # | Dilution (per 2x10^6^ cells) |
| --- | --- | --- | --- | --- |
| Live/Dead | LIVE/DEAD Fixable Yellow Dead Cell Stain | 405/570 | L34959 | 0.2 μL |
| CD86 | Super Bright 436 | 413/431 | 62-0869-42 | 0.25 μg |
| CD206 | Alexa Fluor 488 | 499/520 | 53-2069-42 | 0.0125 μg |
| CD163 | NovaFluor Red 725 | 636/727 | H088T03R05-A | 0.02 μg |

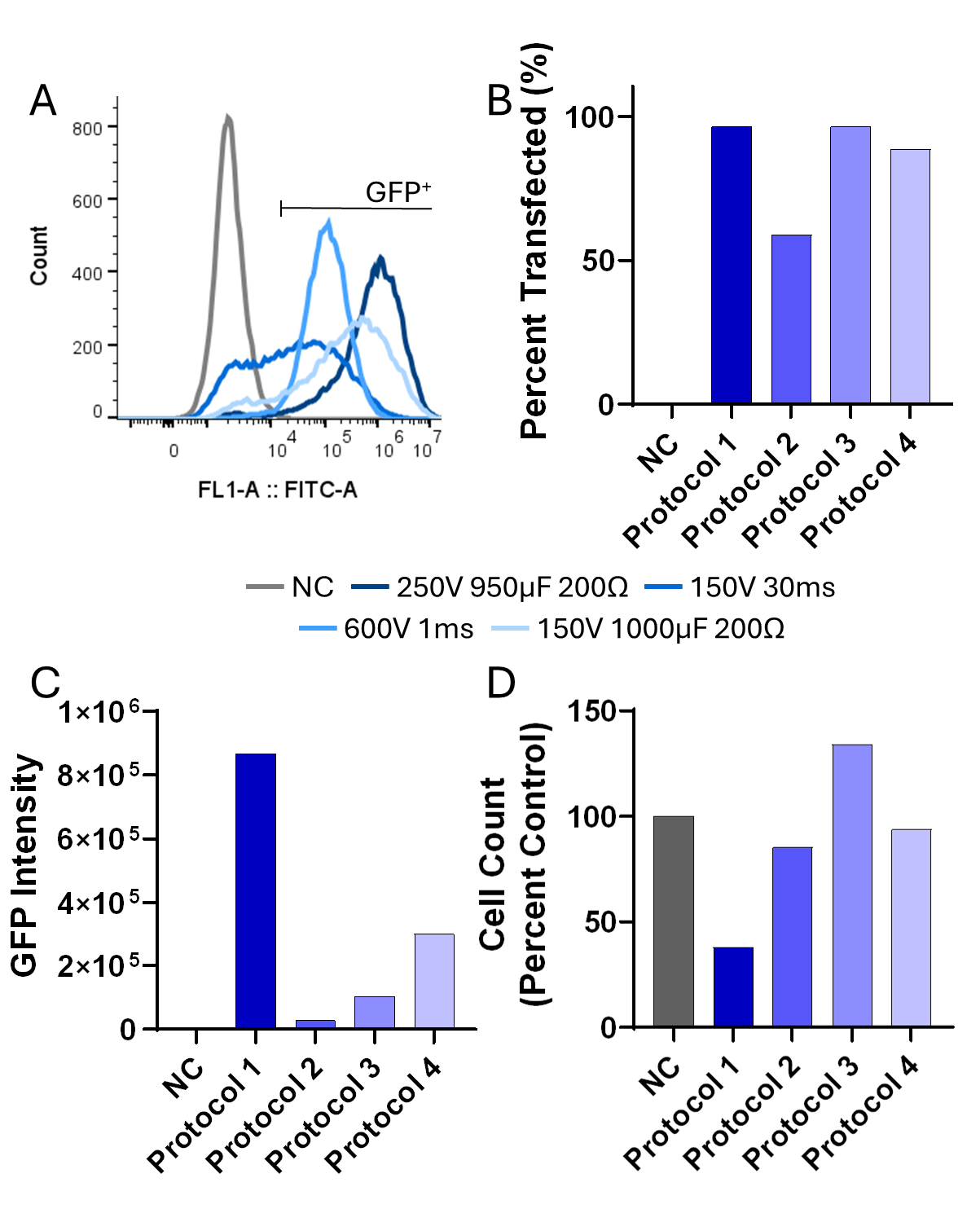

**Figure S1**. Optimization of electroporation protocols. MSCs were transfected using various electroporation parameters to determine the optimal protocol. **A)** Histogram of GFP fluorescent intensities after transfection. Protocol 4 (150 V, 1000 μF, and 200 Ω) was selected based on the **B)** percentage of MSCs transfected, **C)** median fluorescent intensity of the population, and **D)** cell viability represented by the cell count as a percentage of the non-transfected control. Protocol 4 yielded the highest overall number of transfected cells.

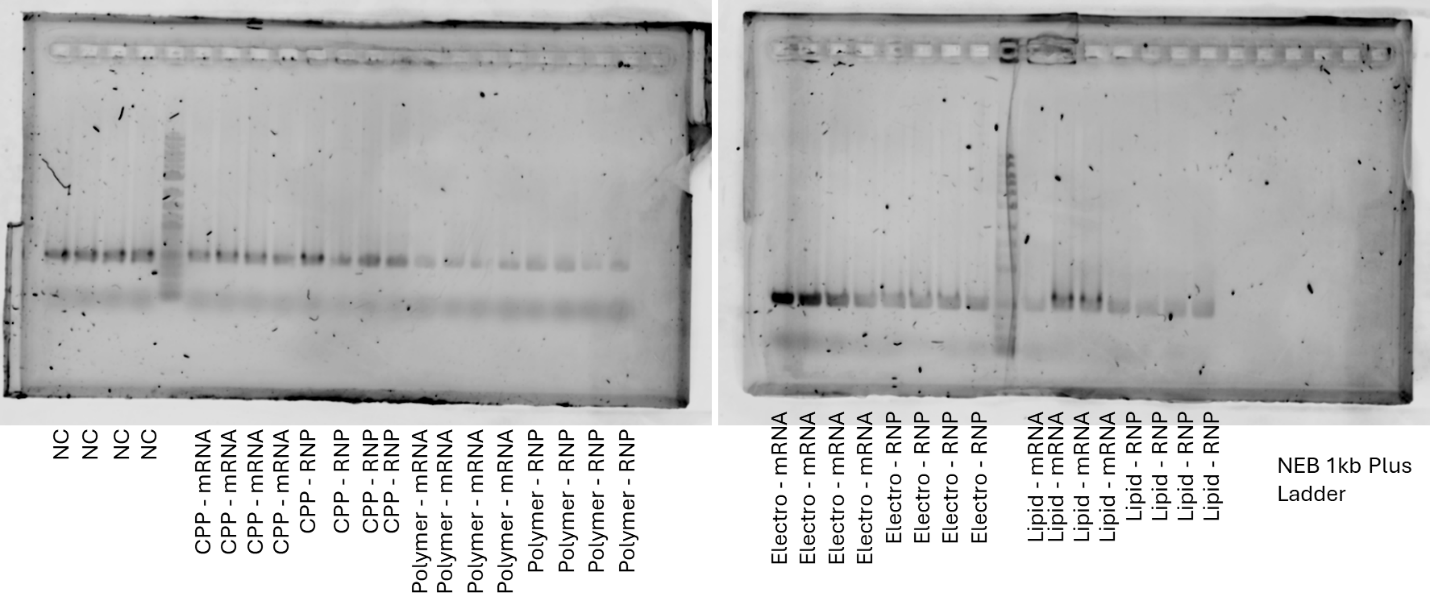

**Figure S2** Amplification of AAVS1 target region using polymerase chain reaction (PCR). Amplified fragments with an expected length of 497 bp were compared to the NEB 1kb Plus DNA ladder.

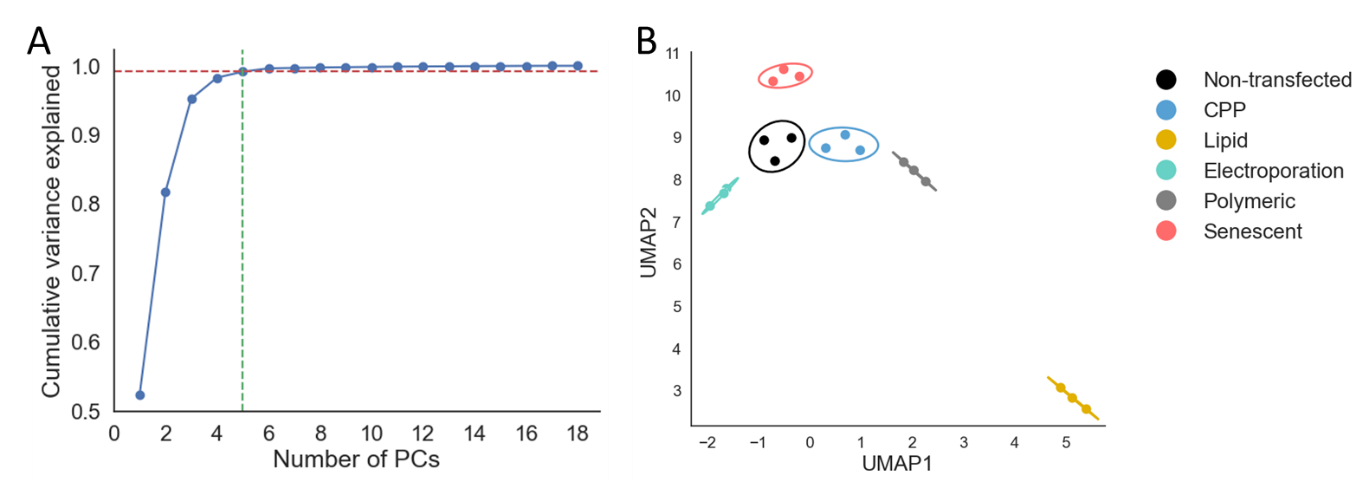

**Figure S3.** UMAP analysis after dimensionality reduction with PCA. **A)** Five principal components (PCs) were selected to maximize the cumulative variance explained without over fitting. **B)** UMAP plot using the top 5 PCs reveals tighter clusters with CPPs and polymers clustered closest to non-transfected cells.

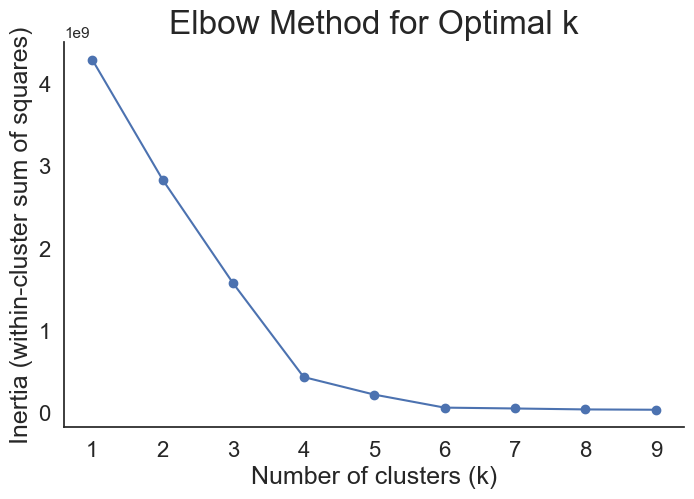

**Figure S4.** Determination of optimal ‘k’ for k-means clustering. At different k values, the Sum of Squared Error (SSE) is calculated between data points and the centroid of their assigned cluster. The optimal k of 5 minimizes the SSE as demonstrated by the ‘elbow’ in the plot.

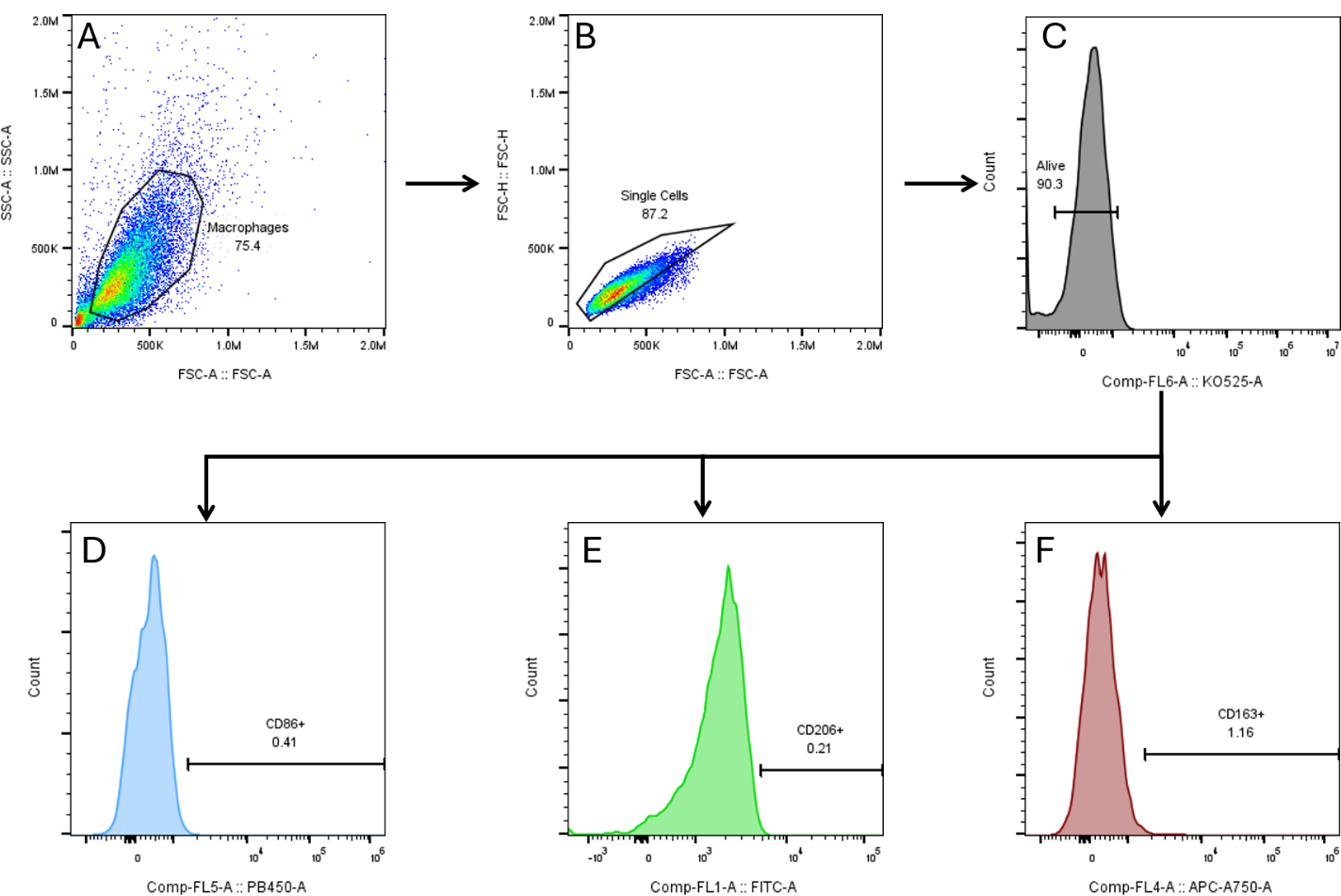

**Figure S5.** Flow cytometry gating strategy. **A)** Macrophages were gated using forward scatter area (FSC-A) versus side scatter area (SSC-A). **B)** Singlets were identified using FSC-height (FSC-H) versus FSC-A. **C)** Live cells were selected by excluding LIVE/DEAD stain-positive cells. Positive populations for each marker were determined using fluorescence minus one (FMO) controls for **D)** CD86 (Pacific Blue), **E)** CD206 (FITC), **F)** CD163 (APC-Cy7). Median fluorescent intensity (MFI) was calculated across the entire live macrophage population.

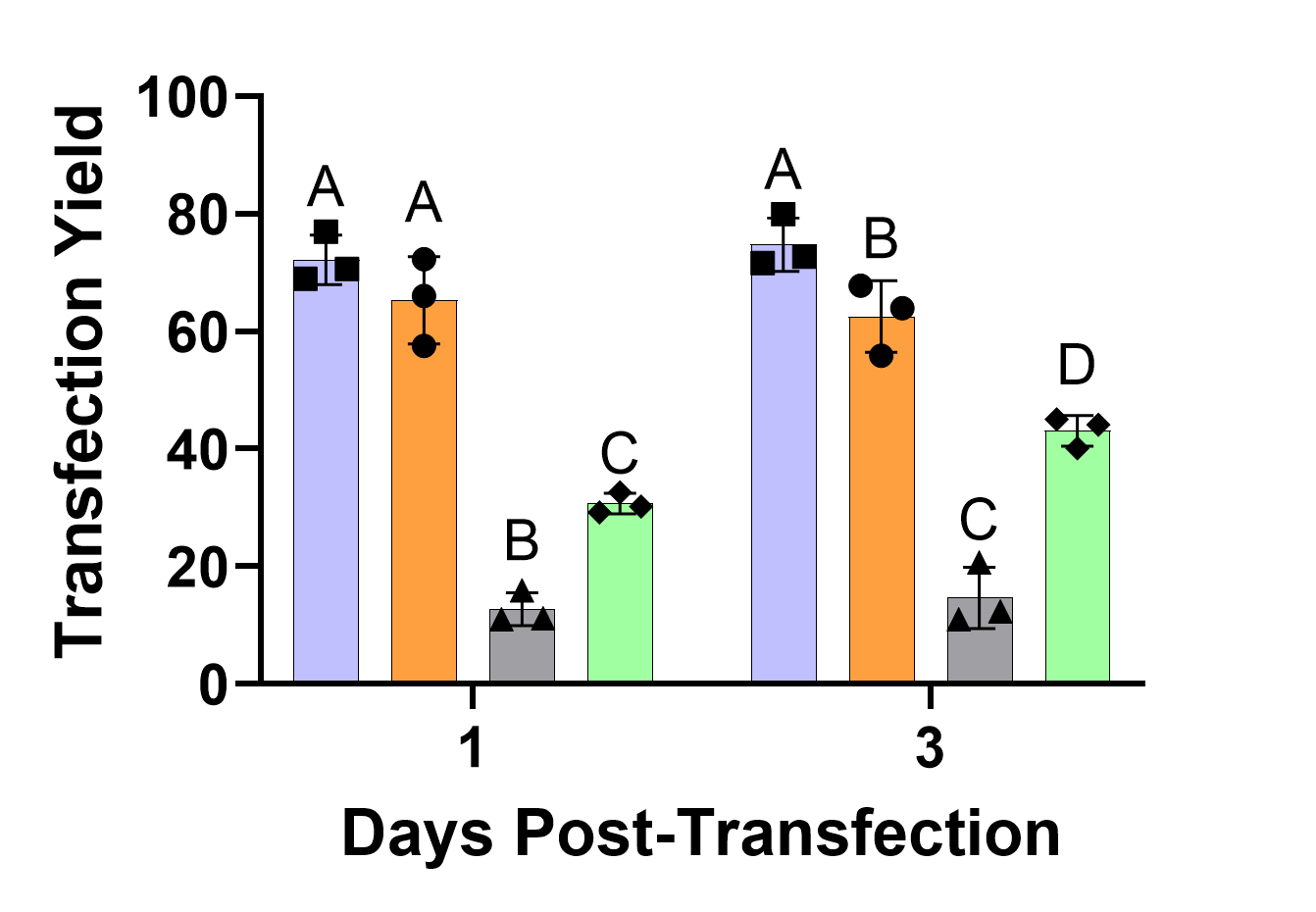

**Figure S6.** Transfection yields from the transfection of eGFP mRNA. The yield incorporates the cell viability and the transfection percentage to give a better representation of the total number of cells expressing the mRNA. Results are plotted as the mean value ± standard deviation for each group (n=3). Statistically similar values are denoted with the same letter.
